# Microglial Fructose Metabolism Is Essential for Glioblastoma Growth

**DOI:** 10.1101/2025.04.12.646414

**Authors:** Leah K. Billingham, Susan L. DeLay, Shashwat Tripathi, Ian Olson, Kaylee Zilinger, Jay Subbiah, Zhaoquan Wang, Nishanth S. Sadagopan, Tzu-yi Chia, Hinda Najem, Guillaume Cognet, Joshua Katz, Gustavo Ignacio Vázquez-Cervantes, Si Wang, Hanxiao Wan, Allie B. Lipshutz, Alina R Murphy, Joeseph Duffy, Irina V. Balyasnikova, Peng Zhang, Dieter Henrik Heiland, Atique U. Ahmed, Catalina Lee-Chang, Amy B. Heimberger, Justin S.A. Perry, Alexander Muir, Navdeep S. Chandel, Jason Miska

**Affiliations:** Department of Neurological Surgery, Feinberg School of Medicine, Northwestern University, 676 N St. Clair, Suite 2210, Chicago, IL 60611, USA; Malnati Brain Tumor Institute of the Lurie Comprehensive Cancer Center, Feinberg School of Medicine, Northwestern University, Chicago IL; Immunology Program, Sloan Kettering Institute, Memorial Sloan Kettering Cancer Center, New York, NY, USA; Immunology and Microbial Pathogenesis Program, Weill Cornell Medicine, New York, NY, USA; Ben May Department for Cancer Research, University of Chicago, Chicago, IL, USA; Department of Neurosurgery, Medical Center – University of Freiburg, Freiburg, Germany; German Cancer Consortium (DKTK), partner site Freiburg, Freiburg, Germany; Department of Neurosurgery, University Hospital Erlangen, Friedrich-Alexander University Erlangen Nuremberg, Erlangen, Germany^3^; Translational Neurosurgery, Friedrich-Alexander University Erlangen Nuremberg, Erlangen, Germany; Louis V. Gerstner Jr. Graduate School of Biomedical Sciences, Memorial Sloan Kettering Cancer Center, New York, NY, USA; Department of Medicine, Northwestern University Feinberg School of Medicine, Chicago, IL, USA

**Keywords:** Brain Tumor, microglia, fructose metabolism

## Abstract

Glioblastoma (GBM) is most common and aggressive primary brain tumor in adults, for which standard of care hasn’t changed in twenty years. GBM tumor associated macrophages (TAMCs), consisting of infiltrating myeloid cells from the periphery and resident microglia cells, are pro-tumorigenic, promoting tumor growth. Fructose is one of the most abundant metabolites in the GBM tumor microenvironment (TME), as well as in the healthy central nervous system (CNS). In the CNS and GBM, microglia are the predominant expressors of the fructose transporter GLUT5. Mice lacking the GLUT5 transporter (GLUT5-KO) survive significantly longer after orthotopic implantation of two glioma cell lines than wildtype mice, which is not due to dietary fructose or peripherally derived TAMCs. Investigation of the TME showed that GLUT5-KO mice have more highly activated and inflammatory innate and adaptive immune compartments. Microglia cultured in fructose have a decreased phagocytic ability and exhibit decreased inflammatory capacity due to the polyol pathway promoting redox homeostasis.

## INTRODUCTION

Glioblastoma (GBM), the most common and aggressive primary CNS cancer in adults, has an extremely poor prognosis of only 16-18 months and nearly 100% fatality. Despite developments of immunotherapies that have proven exceptionally effective in other forms of cancer, the standard of care for GBM, which includes surgical resection, radiation, and temozolomide treatment, only modestly improves patient survival^1^. This is, in part, due to the highly pro-tumorigenic tumor microenvironment (TME), which is characterized by low lymphocyte infiltration and high quantities of immunosuppressive tumor-associated myeloid cells (TAMCs)^2,3^.

TAMCs consist of peripherally derived macrophages and yolk-sac derived brain-resident microglia and play a pivotal role in resistance to both radiation and chemotherapeutic interventions^4–7^ . Recent work by our group and others has identified myeloid cell metabolic phenotypes that promote, or even drive, tumor growth^8–10^. Indeed, modulation of myeloid metabolism in pre-clinical glioma models has been successful at abrogating tumor growth and increasing median survival^9,10^. Much of this work, however, has been done in infiltrating myeloid cells, and the role of microglial metabolism in supporting GBM progression remains unknown.

Microglia exhibit substantial metabolic flexibility, being able to adapt to low-glucose environments in the brain while maintaining functionality by utilizing other carbon sources such as glutamine^11^. Microglia also express the dedicated fructose transporter (GLUT5)^12,13^, which endows them with the ability to metabolize fructose, a metabolite present in high levels in the CNS^14,15^, as a carbon source when glucose availability is low. Despite this and the well-established deleterious effect of high fructose consumption on cardiac, metabolic, and endocrine systems^16,17^, the cellular processes driven by fructose in microglia is unknown. A recent study has demonstrated that a high-fructose diet negatively impacts neurodevelopment in mice via aberrant modulation of microglial function^18^, giving insight into the role of fructose metabolism in immunomodulation in the brain. The question of whether microglia can uptake and metabolize fructose under non-homeostatic conditions—such as in the brain tumor microenvironment—represents an intriguing and entirely unexplored area of study. Using orthotopic glioma models in mice deficient in GLUT5, we tested whether fructose transport by microglia impacts glioblastoma growth. Our data demonstrates that GLUT5 expression in microglia is essential in promoting GBM growth and that its deficiency creates a hyper-inflammatory environment, indicating that targeting fructose metabolism may be powerful target to enhance anti-tumor immunity in GBM.

## RESULTS

### Fructose is abundant in the CNS

To identify the most abundant metabolites in the brain tumor microenvironment (TME), we performed open-flow microperfusion (cOFM)^14,19^ (**Figure 1A**) and in collaboration with the Muir laboratory, performed quantitative metabolomics^20,21^ from the interstitial fluid of CT-2A tumor bearing mice (**Supplementary Table 1)**. Among the most abundant metabolites in the brain were glucose, mannose, and fructose (**Figure 1B**), with the concentration of fructose being nearly half that of glucose (**Figure 1C**, **Supplementary Table 1**). This concentration is surprisingly high compared to plasma but is consistent with previous studies showing that fructose is synthesized in the human brain and is approximately 20-fold higher in cerebrospinal fluid (CSF) than in plasma^14,15^ . Supporting these observations, data obtained from the Human Metabolome Database, shows that while the ratio of CSF glucose to blood glucose is 0.7, the CSF to blood fructose concentration ratio is 5.1 (**Figure 1D**)^22^ . Fructose is primarily transported into cells via the hexose transporter SLC2A5 (GLUT5)^23^ The other known fructose transporter, SLC2A2 (GLUT2) having a substantially higher Km (70-75mM) for fructose than GLUT5 (5-10mM)^24–26^. Analysis of single-cell data from the Protein Atlas^27–30^ demonstrate that GLUT5 is most highly expressed on microglial cells in the human CNS (**Figure 1E**)^27–34^. This suggests that cells of the myeloid lineage might preferentially express GLUT5. However, using the Immunological Genome Project Gene Skyline database^35^ we found that microglia express GLUT5 at levels several log-fold higher than any other immune cell, suggesting a specify to microglia not shared by the rest of the immune system (**Figure 1F**). These data suggest that fructose likely plays a unique and critical role in the CNS and particularly in microglial cells.

**Figure 1.**
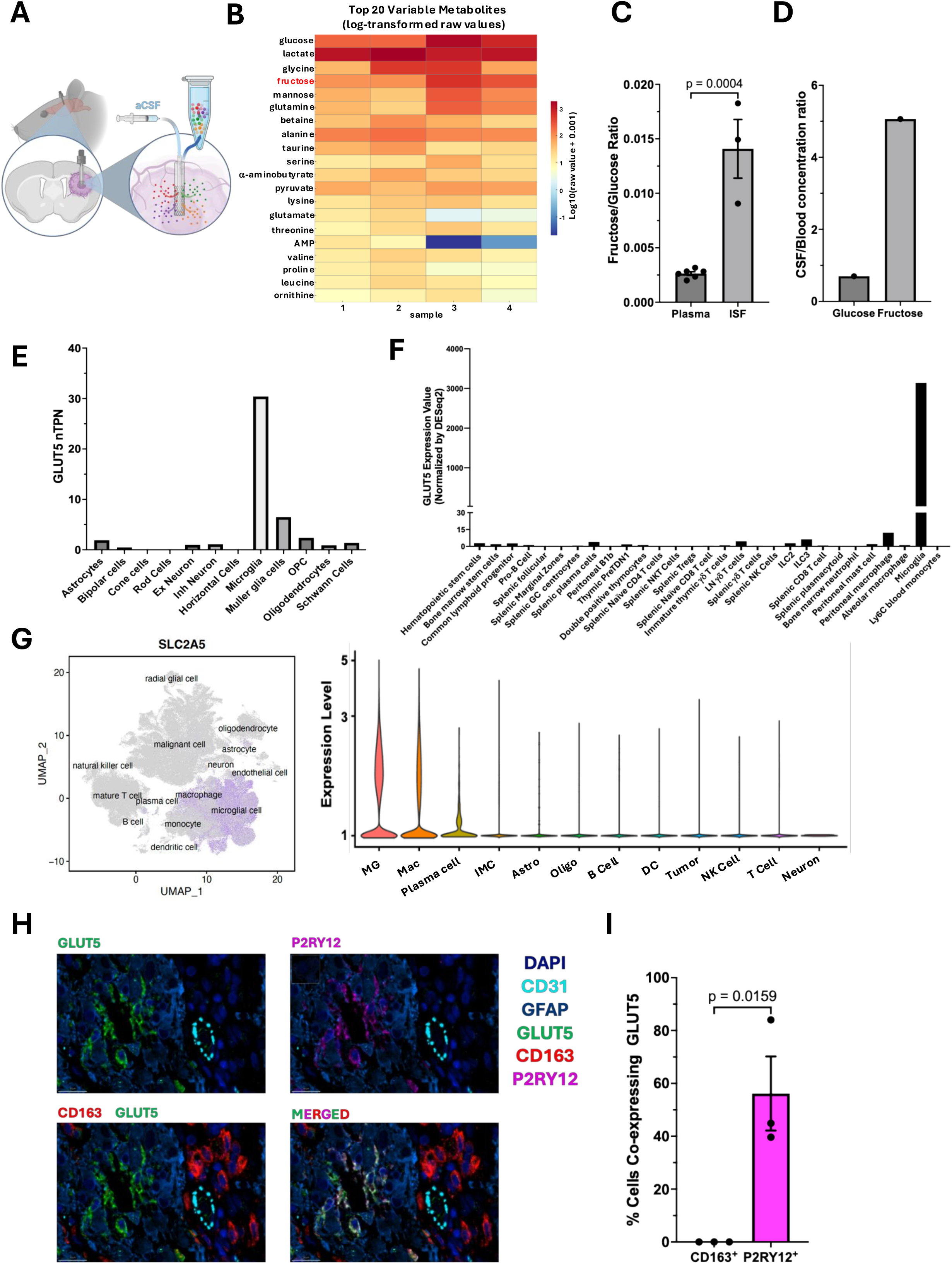
**A.** Schematic of open-flow microperfusion (cOFM) system in mice. **B**. Heatmap of the top 16 metabolites in the interstitial fluid of tumor-bearing murine brains (N=4). **C**. Fructose-to-glucose ratio in plasma and ISF in mice mice (N=6 plasma; N=3 ISF). **D**. Ratio of the average concentration of fructose or glucose in CSF to blood in human samples (Fructose N = avg of 2 reported studies both blood and CSF; Glucose N = avg of 16 reported studies for blood, avg of 3 studies for CSF)^22^. **E**. GLUT5 expression (normalized transcripts per million) in CNS cells from human brains^27–32^. **F**. GLUT5 expression (normalized by DESeq2) in human immune cells^35^ **G**. UMAP (left) of scRNASeq from ∼1 x 10^6 sequenced single cells characterized from 110 GBM patient tumors and level of GLUT5 (right) among GBM TME cells^36^. **H**. Immunofluorescence of GLUT5 (top left); P2RY12 (top right); CD163 and GLUT5 (bottom left), and GLUT5, P2RY12, and CD163 merged (bottom right) in tissue slice from the tumor of a GBM patient (N=3; representative figure shown). **I.** Quantification of data in panel H.

To determine the expression of GLUT5 within the context of CNS malignancy, we analyzed scRNAseq (single-cell RNA sequencing) data from the Gbmap database, which has compiled data from 110 GBM patient tumors^36^. The data indicates that microglial GLUT5 is predominantly expressed in microglia within the tumor microenvironment (TME) while the expression could also be detected in the macrophage cluster and plasma-cells (**Figure 1G)**^36^. Importantly, we could not find any expression by tumor cells. Despite the lack of GLUT5 expression by tumor cells, the expression of GLUT5 is significantly higher in GBM than normal brain tissue (**Extended Data 1A**) and correlated with grade of malignancy in the TCGA database (**Extended Data 1B**).

To further demonstrate the specificity of GLUT5 on microglia in GBM, we performed multiplexed IHC using on newly diagnosed human GBM sections using multiplex immunofluorescencethe with the Lunaphore COMET system. Analysis reveals a striking co-expression of GLUT5 with P2RY12+ (microglia) but not with CD163+ (infiltrating macrophages^37,38^) (**Figure 1H-I).** These findings strongly support that only microglia express GLUT5 in GBM and the broader CNS. This conclusion is reinforced by two key studies. The first^39^ demonstrates that microglial GLUT5 expression is directly regulated by Sall1, a defining transcription factor of the microglial lineage. The second^40^ involves competitive repopulation of the microglial niche following ablation, revealing that only endogenous F4/80^low^ microglia (which distinguished them from F4/80^hi^ bone marrow-derived cells) express both GLUT5 and Sall1.

### Microglia are the only cells in GBM that metabolize fructose

The data above suggests that fructose may only be transported and metabolized by microglia within GBM. To determine if microglia possess a unique ability to metabolize fructose, we cultured CT2A glioma cells, GBM human PDX cells (GBM43), oligodendrocyte cell line MO3.13, RAW 264.7 macrophage cell line, and the BV2 microglial cell line in either glucose-only or fructose-only media for several days and monitored their growth. Surprisingly, while all cell lines were able to proliferate in glucose-only media, only the BV2 microglia were able to proliferate in fructose-only media (**Figure 2A)**. Moreover, only the BV2 cells were able to survive at all in fructose alone (**Figure 2B**). This data demonstrates that microglia are programmed to utilize fructose for survival where other cells of the brain tumor TME cannot.

**Figure 2.**
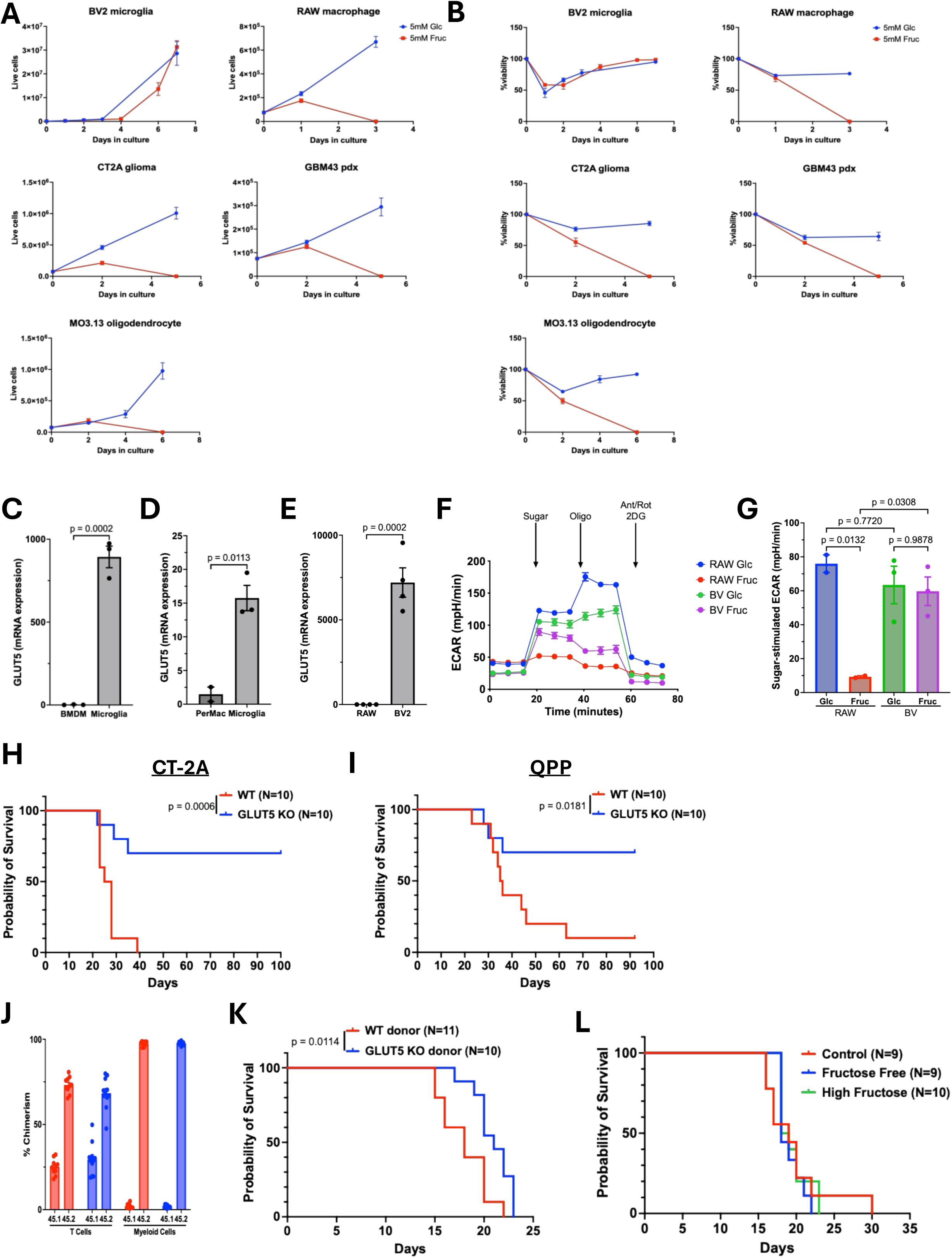
**A.** Proliferation of BV2 microglia, RAW macrophages, CT2A glioma, GBM43 human PDX, and MO3.13 oligodendrocyte cell lines in media containing either 5mM glucose or 5mM fructose (N=3). **B**. Viability of BV2 microglia, RAW macrophages, CT2A glioma, GBM43 human PDX, and MO3.13 oligodendrocyte cell lines in media containing either 5mM glucose or 5mM fructose (N=3). **C-D**. GLUT5 mRNA expression in microglia relative to (C) BMDMs, and (D) peritoneal macrophages and microglia isolated from adult mice (N=3 BMDM, Microglia; N=2 Permac). **E**. GLUT5 mRNA expression in RAW macrophages and BV2 microglia (N=4 RAW macrophages; N=3 BV2) **F.** Extracellular acidification rate of RAW or BV2 cells fed 5mM glucose or 5mM fructose (N=2 RAW, N=3 BV2; representative figure shown). **G**. Sugar stimulated ECAR in RAW or BV2 cells fed with 5mM glucose or 5mM fructose (N=2 RAW, N=3 BV2) **H-I**. Kaplan Meier curves of mice implanted i.c. with 1×10^5 CT-2A or QPP4 glioma cells (respectively) (A:N=10; B: N=10). **J**. Percent of CD45.1 and CD45.2 myeloid and T cells in peripheral blood of CD45.1 mice 6 weeks post irradiation and transplant of bone marrow from CD45.2 WT and GLUT5 KO mice. **K**. Kaplan Meier curves of recipient CD45.1 mice transplanted with 1e6 bone marrow cells from either WT or GLUT5KO recipients and subsequently implanted with 100K CT-2A glioma cells (N=11 CD45.1 with WT donor; N=10 CD45.1 with GLUT5 KO donor). **L.** Kaplan Meier curves of C57Bl/6 mice fed either a control diet, a fructose-free diet, or a high-fructose (60%) diet and subsequently implanted with 100K CT-2A glioma cells (N=9 control feed; N=9 fructose-free feed; N=10 high-fructose feed).

While this data supports that only microglia can metabolize fructose, RAW macrophages and BV2 are indeed cell lines, so we sought to validate the GLUT5 expression in primary microglia, peritoneal macrophages, and bone marrow-derived macrophages from adult WT mice. The microglia expressed 15 times more GLUT5 than peritoneal macrophages (Permac) (p=0.0113) and approximately 900 times more than bone marrow-derived macrophages (BMDMs) (p=0.0002) (**Figure 2C-D)**. Similarly, the microglial cell line BV2 around 7,000 times more GLUT5 than theRAW macrophage cell line (p=0.0002) (**Figure 2E)**. To assess whether fructose is actively metabolized, we measured the extracellular acidification rate (ECAR) of RAW macrophages and BV2 microglia using a Seahorse extracellular flux analyzer following treatment with 5 mM glucose or fructose. The glucose-stimulated ECAR was ∼76mpH/min in RAW macrophages and ∼63mpH/min in BV2 microglia, which was not significantly different (p=0.7720). However, the fructose-stimulated ECAR in RAW macrophages was only ∼9mpH/min compared to ∼60mpH/min in the BV2 microglia. Importantly, there was no difference sugar-stimulated ECAR when fed glucose compared to fructose in BV2 microglia (p=0.9878). Thus, RAW macrophages were unable to metabolize fructose whereas BV2 microglia could (**Figure 2F-G**). Notably, both RAW and BV2 cells were able to maintain oxygen consumption regardless of which sugar they metabolized (**Extended Figure 2A-B**). We were not able to successfully perform seahorse analysis on primary microglia.

To determine if fructose transport by microglia influences brain tumor growth, we implanted the glioma cell line CT2A into WT mice and mice globally deficient in GLUT5 (GLUT5-KO mice). Two weeks after tumor implantation, the CD45+ immune cell populations in the tumor were analyzed using scRNASeq (**Extended Figure 3A)**. Tumors from GLUT5-KO mice had no expression of SLC2A5 (**Extended Figure 3B**). In wild-type mice, P2ry12-expressing microglia—but not peripherally derived CD49D^+^ myeloid cells^38^—expressed GLUT5 within the tumor microenvironment (**Extended Figure 3C-D**).

### GLUT5 deficiency induces rejection of orthotopic brain tumors

As fructose is highly abundant in the CNS and the transporter is expressed by microglia both in tumor-bearing and healthy brain tissue (**Figure 1)**, we hypothesized that fructose metabolism in the brain may play a role in glioma tumor growth. We implanted the immunotherapy-resistant CT2A glioma cell line^41^ or QPP4 gliomas derived from the Qki genetically engineered murine model^42,43^ into WT and GLUT5-KO mice. Surprisingly, 70% of GLUT5-KO mice implanted with either cell line did not succumb to disease, whereas all WT mice implanted with CT-2A tumor cells and 90% of mice implanted with QPP4 died (**Figure 2H-I)**, demonstrating that fructose transport in the host is critical for glioblastoma growth. GLUT5KO mice consistently outlived WT across all three experiments with CT-2A tumors, although survival percentages varied (**Extended Data 4**). Given our own data and other recent studies^44,45,40^ that indicate GLUT5 expression in the brain is specific to microglia, we generated bone marrow chimera mice by i.v. injecting the bone marrow from WT or GLUT5-KO mice into irradiated CD45.1 C57Bl/6 mice. After allowing the immune compartment to repopulate for 8 weeks (**Figure 2J**), we implanted CT-2A cells and monitored mice for survival. While there was a modest survival benefit in the mice that received bone marrow from GLUT5-KO mice (**Figure 2K**), this survival benefit was not comparable to the global GLUT5-KO implanted with CT-2A, indicating that the bone marrow-derived myeloid cells do not drive the survival benefit.

Beyond microglia in the brain, GLUT5 is primarily expressed at high levels in the small intestines, liver, testes, and sperm^34,46^ . As the GLUT5 is globally knocked out, we considered that the inability of mice to uptake fructose from the diet could be driving the robust survival of the GLUT5-KO mice after tumor implantation and that a high-fructose diet would decrease median survival while a fructose-free diet would increase it. To address this, we fed WT mice a control diet, fructose-free diet, or a high-fructose (60%) diet for two weeks prior to implantation of CT2A tumor cells. Interestingly, neither a fructose-free diet nor a high-fructose diet had any impact on median survival (**Figure 2L**), indicating that dietary fructose is not responsible for the observed phenotype.

### GLUT5 deficiency leads to hyperinflammatory immune phenotypes in the CNS

To understand the process underlying tumor rejection in GLUT5-KO mice, we utilized our scRNASeq data to assess the immune compartment of tumors from WT and GLUT5-KO mice 14 days post-implantation of CT2A cells (**Extended Figure 5**). Analyses revealed that the immune compartment of the TME in GLUT5-KO mice was markedly more activated than that from WT mice (**Figure 3A-B)**. Microglia from GLUT5-KO mice exhibited higher expression of genes critical for antigen presentation (H2-K, CD274, H2-K1, H2-D1, H2-Ab1) and inflammation (Stat1, IFNgr1) (**Figure 3A)**. Similarly, CD8+ T cells from GLUT5-KO mice had increased expression of activation markers (CD44, Stat3, Stat1, etc) as well as anti-tumor mediators (Stat1, Gzmb) (**Figure 3B**). Interestingly, GSEA analysis uncovered an significant enrichment in IFNg response genes in GLUT5-KO microglia (**Figure 3C).** Furthermore, network interactome analysis^47^ revealed more interaction of microglia with effector T cell populations (**Figure 3D-E**) and more interaction of activated CD8+ T cells overall (**Figure 3F**) in GLUT5-KO mice. These indicate that GLUT5 expression in microglia controls immune activation and effector activity in both the innate and adaptive immune compartments, which is likely dependent on the interaction of microglia with adaptive immune cells.

**Figure 3.**
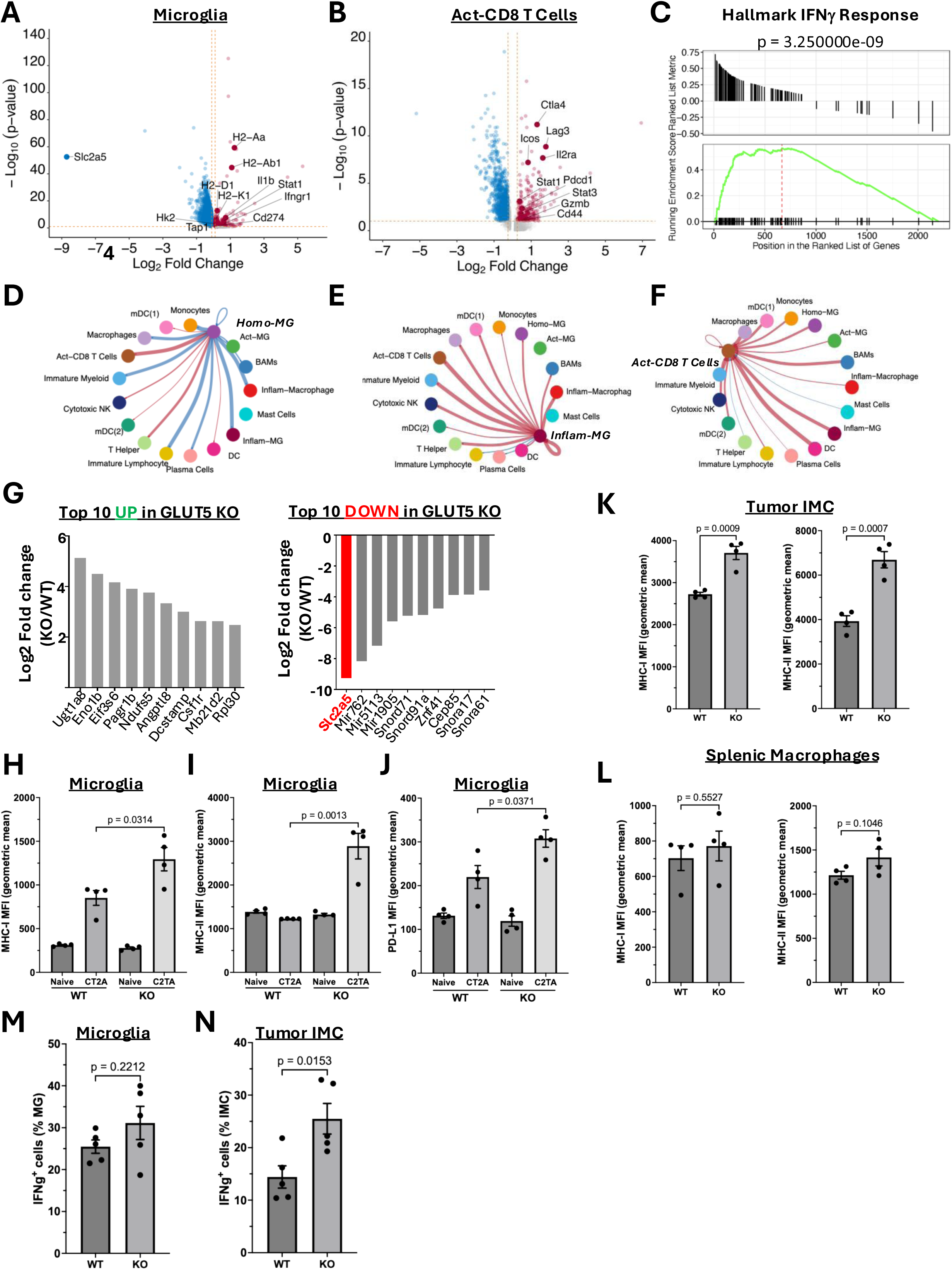
**A.** Volcano plot of differentially expressed genes in microglia from scRNASeq of CD45+ cells in CT-2A glioma cell tumors (N=1 each genotype). **B.** Volcano plot of differentially expressed genes in activated CD8+ T cells from scRNASeq of CD45+ cells in CT-2A glioma cell tumors (N=1 each genotype). **C.** GSEA of hallmark interferon gamma response across all CD45+ cells from CT=2A glioma cell tumors(N=1 each genotype). **D-F**. Strength of cell-cell interactions of homeostatic microglia (D), inflammatory microglia (E)and activated CD8 T cells (F) with other CD45+ cell populations in CT-2A glioma cell tumors from KO mice compared to those from WT mice. Red indicates increased levels of interaction; blue indicates decreased levels of interaction. (N=1 each genotype). **G**. Top 10 upregulated (left) and upregulated (right) genes in microglia isolated from naïve adult GLUT5 KO mice relative to WT mice (N=4 WT; N=3 GLUT5-KO). **H-J**. MFI (geometric mean) of MHC-I (H), MHC-II (I), and PDL-1 (J) on microglia of naïve mouse brains or 2 weeks after implantation with 100K CT-2A glioma cells (N=4 each group). **K.** MFI (geometric mean) of MHC-I (left) and MHC-II (right) on infiltrating myeloid cells to tumor-bearing brains (N=4 each group). **L.** MFI (geometric mean) of MHC-I (left) and MHC-II (right) on splenic myeloid cells from tumor-bearing mice 2 weeks after implantation with 100K CT-2A glioma cells (N=4 each group). **M-N**. Percent IFNg+ microglia (M), infiltrating myeloid cells (N) fromvbrains of tumor-bearing mice 2 weeks after implantation with 100K CT-2A glioma cells (N=4 each group)

To determine whether resting microglia from GLUT5-KO mice were more inflammatory at baseline, we performed bulk RNASeq on microglia from naïve WT and GLUT5-KO brains. Several of the top upregulated genes such as UGT1A8, Eno1b, and Angptl8 are involved in metabolism (**Figure 3F-G**). One of the genes, Mb21d2, plays a positive role in modulating the cGAS-STING pathway in myeloid cells and deficiency leads to impaired production of type 1 interferon^48^ . None of the downregulated genes indicated changes in basal levels of inflammation (**Figure 3G**). These data suggest that the increase in inflammation observed in the glioma TME of GLUT5-KO brains is in response to a stimulus rather than a steady-state/developmental phenomenon.

To determine whether there were any baseline immunological differences in microglia in GLUT5 knockout (KO) mice, we performed immunophenotyping on naïve and tumor-bearing brains from WT and GLUT5-KO mice implanted with CT2A glioma cells. No differences were observed in the cell surface expression of MHC-I, MHC-II, or PDL-1 between WT and GLUT5-KO microglia from healthy brains (**Figure 3H–J**). Conversely, tumor-bearing brains from GLUT5-KO mice implanted with CT2A cells exhibited significantly higher levels of MHC-I, MHC-II, and PDL-1 on the cell surface of microglia compared to WT controls (p=0.0314, p=0.0013, and p=0.0371, respectively). These data demonstrate that, in the presence of glioma, microglia unable to uptake fructose become more highly activated. Tumor-infiltrating myeloid cells also expressed higher levels of MHC-I and MHC-II in GLUT5-KO mice relative to WT (**Figure 3K)**. Conversely, there was no difference in expression of MHC-I, MHC-I, in the spleens between WT and GLUT5-KO mice (**Figure 3L**) indicating that the effect of GLUT5-KO is preferential to the brain compartment and that changes in microglial metabolism drive anti-tumor immune responses. We furthermore observed increased IFNg expression in infiltrating myeloid cells as well as an upward trend in microglia (**Figure 3M**).

### GLUT5 deficiency leads to enhanced adaptive immunity against GBM

In the TME of GLUT5-KO mice, CD8+ T cells were more highly activated as observed by increased expression of IFNg (**Figure 4A**) and Granzyme B (**Figure 4B**). To determine whether the increase activation of CD8+ T cells was antigen specific, we implanted WT and GLUT5-KO mice with CT2A-OVA glioma cells, which express the ovalbumin (OVA) protein^49^ . There was nearly double the percentage of OVA-specific CD8+ T cells in the gliomas of the GLUT5-KO mice compared to the WT (p=0.03), indicating that CD8+ T cells in this TME were adept at responding to a specific antigen (**Figure 4C).** Supportive of this, scRNASeq data revealed that T cells from GLUT5 KO tumor bearing mice had a decreased percent of unique clones (**Figure 4D**) and that the top 20 clones in the GLUT5-KO TCR repertoire make up a greater percentage of total TCR clones than the top 20 in the WT (**Figure 4E**). These data suggest increased clonal expansion of clones, allowing for a more targeted antitumor response.

**Figure 4.**
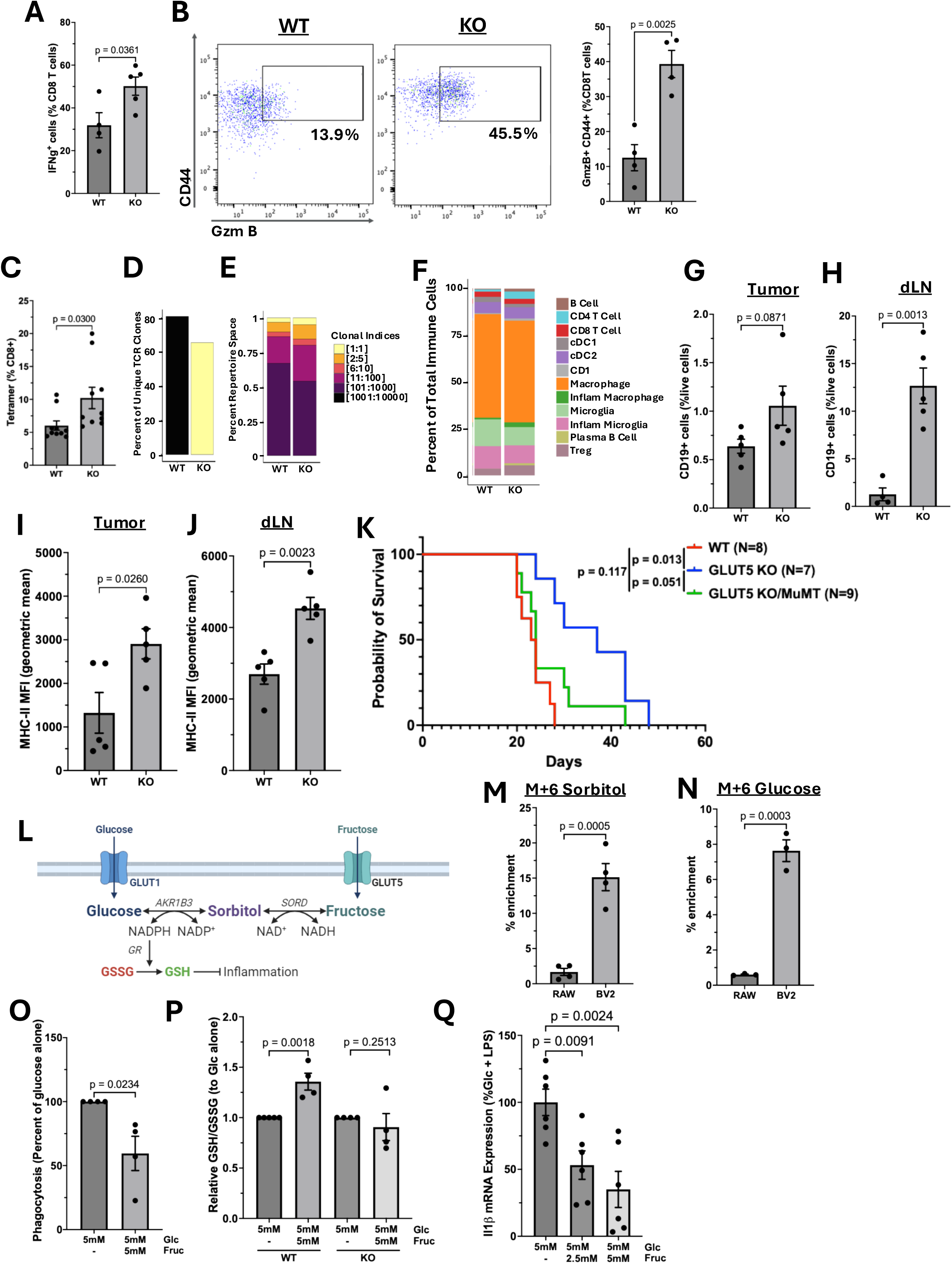
**A.** Percent IFNg+ CD8 T cells from brains of tumor-bearing mice 2 weeks after implantation with 100K CT-2A glioma cells (N=4 each group). **B.** Representative figure (left) and percent GzmB+ (right) CD8+/CD44+ T cells in tumor-bearing brains (N=4). **C**. Percent tetramer positive CD8 T cells in tumor bearing brains 2 or 3 weeks after implantation with CT-2A glioma cells (N=10 both groups). **D**. Percent of unique TCR clones in WT vs KO T cells. **E**. Size of clonal indices in WT vs KO T cells. **F**. Immune cell composition of tumors from WT vs KO tumor-bearing mice (N=1). **G-H**. Percent B cells of live cells in brains (G) and draining lymph nodes (H) from tumor-bearing WT vs KO mice (N=4 WT, N=5 KO). **I-J**. MFI (geometric mean) of MHC-II on B cells in the brains (I) or lymph nodes (J) of tumor-bearing WT or KO mice (N=5 WT, N=4 KO). **K.** Kaplan Meir curves of WT, GLUT5 KO, or GLUT5 KO/MuMT mice after implantation with 100K CT-2A glioma cells (N=8 WT, N=7 GLUT KO, N=9 GLUT KO/MuMT). **L**. Schematic of the polyol pathway. **M**. Percent enrichment of M+6 13C-labeled sorbitol in RAW macrophages and BV2 microglia after 8h of culture with 13C-Fructose. **N**. Percent enrichment of M+6 13C-labeled glucose in RAW macrophages and BV2 microglia after 24h of culture with 13C-Fructose. **O**. Percent phagocytosis of apoptotic CT-2A glioma cells by primary microglia cultured in either 5mM glucose or 5mM glucose and 5mM fructose. Data presented as the percent phagocytosis of the glucose only samples treated on the same day (N=4 from 2 individual experiments). **P.** Relative GSH/GSSG ratio of LPS-treated primary microglia from WT and GLUT5-KO mice cultured with either 5mM glucose or 5mM glucose and 5mM fructose. Data represented as the ratio in Glucose/Fructose cultured cells relative to microglia culture in glucose alone. (WT, N=4 from 3 individual experiments; KO, N=4 from 2 individual experiments). **Q**. Il1b mRNA expression in BV2 cells treated with 100ng/mL LPS for 4hin the presence of 5mM glucose, 5mM glucose + 2.5mM fructose, or 5mM glucose + 5mM fructose. Data presented as the percent mRNA fold change expression of the averaged glucose only samples treated on the same day (N=6 from 3 individual experiments).

Interestingly, our scRNASeq data revealed that tumors from GLUT5 KO mice have a greater abundance of B cells and plasma B cells (**Figure 4F**), suggesting that B cells may play a critical role in our observed phenotype. Flow cytometric analysis of the brains (**Figure 4G**) and draining cervical lymph nodes (**Figure 4H**) from these mice further confirmed increased B cell infiltration, as evidenced by CD19 expression. Moreover, B cells in both the brain (**Figure 4I**) and lymph nodes (**Figure 4J**) expressed elevated levels of MHC-II. To assess whether B cells play a pivotal role in the rejection of tumors in GLUT5 KO mice, we crossed GLUT5 KO mice with mice that do not have B cells (muMT). We then implanted WT, GLUT5 KO, and GLUT5 KO/MuMT mice with the CT-2A glioma cell line and monitored survival. While GLUT5 KO mice had an increased median survival compared to WT mice, the absence of the fructose transporter did not offer any survival benefit in the absence of B cells (**Figure 4K**). Thus, B cells are critical to the anti-tumorigenic immune microenvironment driven by the absence of microglial fructose metabolism. Overall, these data demonstrate that the lack of fructose metabolism in microglia promotes a more highly activated adaptive immune responses against GBM.

### Fructose restricts inflammatory activation of microglia by promoting redox homeostasis

A recent study reported that fructose suppressed the polarization of TAMs into M1-like cells and that this led to increased tumor size in multiple models of colorectal cancer^50^ . Since impaired fructose uptake in microglia results in an enhanced inflammatory phenotype, both at the cellular level and within the tumor microenvironment, we sought to determine whether the presence of fructose could attenuate the inflammatory response in microglia. We co-cultured primary microglia isolated from neonate mice (p1-p4) with apoptotic CT2A glioma cells in either media containing 5mM glucose or 5mM fructose and 5mM glucose and measured the percent of phagocytic microglia. Microglia that had been cultured with fructose were less phagocytic than those cultured in glucose alone (**Figure 4O**). Phagocytosis is a critical step in antigen presentation, and blockade of phagocytosis by fructose could lead to a decreased ability of microglia to clear tumor cells and present tumor antigens to promote antitumor functionality.

It is well established that in both microglia and macrophage inflammatory activation is typically paired with the upregulation of reactive oxygen species^51–55^. Importantly, in the context of diabetic neuropathy, it is thought that the conversion of glucose to fructose produces inflammation through the polyol pathway which can deplete NADPH and NAD+ which prevents the ability to handle ROS^56^. Therefore, we hypothesized that if the polyol pathway ran in reverse (i.e. converted fructose to glucose) you could regenerate NADPH and promote redox homeostasis (**Figure 4L**). We performed ^13^C-fructose flux tracing in RAW macrophages and BV2 microglia. After 8h, we observed 15% enrichment of M+6 sorbitol in BV2 microglia (**Figure 4M)**, and 7.5% enrichment of M+6 glucose in BV2 microglia after 24h (**Figure 4N**). Importantly, there was no incorporation of labeled fructose into sorbitol or glucose in RAW macrophages. This demonstrates that microglia can produce glucose from fructose.

The conversion of fructose to glucose via the polyol pathway produces NADPH, an electron donor essential for maintaining cellular redox balance. NADPH is critical for the recyling of glutathione from its oxidized form (GSSG) to its reduced form (GSH)^57^. We observed that primary microglia exposed to LPS for 4h had a higher GSH/GSSG ratio in the presence of fructose compared to glucose alone (**Figure 4P**). This suggests that fructose supports increased recycling of GSSG to GSH, which could mitigate ROS-induced inflammation. Notably, we did not see an increase in the GSH/GSSG ratio in the presence of fructose in microglia harvested from GLUT5-KO mice, indicating that this difference is due to the presence of fructose in the microglia.

To determine whether the presence of fructose decreases the inflammatory response, BV2 microglial cells were cultured in media containing 5 mM glucose, 5 mM glucose + 2.5 mM fructose, or 5 mM glucose + 5 mM fructose. After a 4-hour LPS stimulation, we measured the expression of the pro-inflammatory cytokine Il1b. Notably, the addition of 2.5mM fructose significantly reduced IL-1b expression by 50% (p=0.0091), and 5mM of fructose reduced Il-1b expression by 70% (p=0.0024) (**Figure 4Q**). These data point to a role for fructose to dampen the inflammatory response in microglia, potentially through the sustained antioxidant effect of increased GSH.

## DISCUSSION

Here we show that in the GBM tumor microenvironment, microglia are the sole expressors of the fructose transporter GLUT5. Furthermore, we demonstrate that fructose uptake and metabolism by microglia supports the progression of GBM through the inhibition of inflammation via modulation of GSH levels. GLUT5 KO mice have enhanced survival and a more highly activated GBM microenvironment characterized by increased antigen presentation by infiltrating myeloid cells, microglia, and B cells, increased TCR specificity, higher IFNg production and response, and increased infiltration of B cells.

The finding that, in the context of GBM, fructose metabolism could promote an anti-inflammatory on the surface seems to contradict the myriad of studies linking fructose consumption to lipogenesis, obesity, metabolic syndrome, cancer progression, and inflammation^16,58–60^. However, as the survival benefit observed in GLUT5 KO mice could not be recapitulated with a fructose free diet (**Figure 2**) and the CNS is capable of producing its own fructose via the polyol pathway^61,62^, it is possible that microglial fructose metabolism is independent of fructose elsewhere in the body. Other work citing fructose as a necessary metabolite to promote cancer growth^63,64^ has been done in cells that produce their own fructose from glucose via the polyol pathway. It’s important to note that this is due to an abundance of glucose rather than an increased uptake of fructose, and thus likely has different biological effects. Furthermore, as the CSF has a vastly higher fructose-to-glucose ratio than plasma (**Figure 1**), it’s possible that microglia have adapted to utilize fructose in a manner that promotes homeostasis rather than disrupts normal tissue function. An anti-inflammatory effect of fructose, rather than an inflammatory effect, on microglia could help mitigate inflammation that could cause brain injury upon assault. Whether a lack of GLUT5 in other models of brain injury, such as stroke or traumatic brain injury, remains to be seen.

While our models, and the work of others^40,65^, showed that fructose uptake and metabolism is unique to microglia among immune cells, some studies have shown that peripheral macrophages and monocytes can express the transporter *in vitro* at low levels ^4^/12/2025 9:29:00 PM One recent study showed uptake of fructose by WT BMDMs cultured in M38-conditioned media was greater than that of BMDMs from Glut5fl/fl × Lyz2-cre mice.

Furthermore, they showed that the Glut5fl/fl × Lyz2-cre mice had slower growing tumors in a colorectal cancer model^50^, suggesting that metabolism of fructose by TAMs could be shaped by the specific tumor microenvironment in different tissues to promote tumor growth. The impact of fructose metabolism is, indeed, likely dependent on cell type. In another recent study, genetic engineering of CD8 T cells to express GLUT5 leads to higher anti-tumor immunity^66^ . This is an intriguing area of study given the diversity of immune infiltration in different tumors and how the inflammatory or anti-inflammatory action of fructose among tumor infiltrating immune cells is dictated both by the TME and the cell type metabolizing fructose.

Our study highlights the importance of microglia and fructose metabolism in GBM, something that has been minimally explored. Whether the impact of fructose metabolism on microglia function impacts initial tumorigenesis or the later anti-tumor response is an intriguing question not addressed in this study, especially given recent evidence of a role for microglia in gliomagenesis^67^ . More work needs to be done to assess how microglial fructose uptake could be altered *in vivo* to promote anti-tumor immune cell function in GBM.

## MATERIALS AND METHODS

### GLIOMA CELL LINES AND TUMOR IMPLANTATION

The CT-2A tumor line was initially obtained from Sigma/Millipore. CT-2As were cultured in Dulbecco’s modified Eagle’s medium (Corning) supplemented with 10% fetal bovine serum (FBS; HyClone), 1% penicillin-streptomycin, and glutamax. QPP4 cells were derived from Nestin-CreERT2 QkL/L; Trp53L/L; PtenL/L mice^42,43^ and kindly provided by Dr. Amy B Heimberger (Northwestern University). They were culture in suspension with DMEM/F12 supplemented with 1% penicillin/streptomycin, 2% B27, 10 ng/mL EGF, and 20 ng/mL FGF. Prior to implantation, CT-2A were lifted with trypsin-EDTA (Corning). QPP4 were broken up with accutase. All cells were washed with phosphate-buffered saline and suspended in PBS at a concentration of 40 x 106 cells/mL. 1 x 105 cells were injected intracranial at a 3-mm depth using a stereotactic apparatus.

### CELL CULTURE

BV2 microglia cells were obtained Accegen. Human PDX GBM43 cells were obtained from David James (Northwestern University). RAW macrophages were obtained from ATCC. BV2 were cultured in sugar free DMEM [Fisher; 11-966-025] supplemented with 10% FBS, 1% penicillin-streptomycin, glutamax and either 5mM glucose, 5mM glucose/2.5mM fructose, or 5mM glucose/5mM fructose as indicated in figure legends. CT-2A, human PDX GBM43, MO3.13, and RAW cells were cultured in Dulbecco’s modified Eagle’s medium (Corning) supplemented with 10% fetal bovine serum (FBS; HyClone), 1% penicillin-streptomycin, and glutamax.

### FRUCTOSE FLUX

DMEM was supplemented with 10% FBS, 1% Penicillin/Streptomycin, 5mM glucose, and 2.5mM U13C-fructose. BV2 and RAW cells were washed with PBS prior to treatment with 13C media for 8 and 24h. Media was aspirated and cells were washed 2x with ice-cold saline before scraping into 1mL ice-cold 80% methanol/20% H2O overnight. Samples were lysed by 3 freeze/thaw cycles using liquid nitrogen followed by 42C water bath; after freeze/thaw, samples were spun at 16,900 xg for 15m. Supernatant was collected and samples were analyzed by High-Performance Liquid Chromatography and High-Resolution Mass Spectrometry and Tandem Mass Spectrometry (HPLC-MS/MS) as previously described^9^.

### PERITONEAL MACROPHAGE ISOLATION

Peritoneal cells were collected via peritoneal lavage with 5mL cold PBS. Lavage was spun for 5mL at 300xg and resuspended in serum-free RPMI before plating in a 6-well plate. Macrophages were allowed to adhere for 45 minutes, at which point cells were washed 2x with PBS and collected for RNA isolation.

### BMDM ISOLATION AND CULTURE

Bone marrow was isolated from C57BL/6J mice and subjected to red blood cell lysis with ACK lysis buffer prior to neutralization with PBS and resupsension in RPMI supplemented with 10% FBS, 1% penicillin/streptomycin, and 20ng/mL mCSF (Thermo Fisher AF-315-02).

Cells were counted using a hemocytometer and plated in 10cm tissue culture plates at a density of 3-million cells/plate. Cells were cultured with 20ng/mL mCSF to induce differentiation into BMDMs. Media was changed every 3 days and BMDMs were harvested by scraping on day 6 and collected for RNA isolation.

### CELL PROLIFERATION

BV2 murine microglia, GBM43 PDX human glioma, CT2A murine glioma, MO3.13 human oligodendrocytes, and RAW murine macrophages were each cultured in sugar-free DMEM supplemented with 10% fetal bovine serum, 1% penicillin/streptomycin, 1% glutaMAX, 1% sodium pyruvate, 5mM glucose and 2.5mM fructose for 3 days before being transferred into 12-well plates, 75,000 cells per well with media containing either 5mM glucose or 5mM fructose. Cells were allowed to grow to confluence, counted with trypan blue staining, and expanded to increasing plate sizes. Samples that did not grow to confluence or died during culture were not counted. Proliferation was monitored until all cells in fructose-only media had died.

### FRUCTOSE-MODIFIED DIET

Custom mouse feed was purchased from TestDiet. Mice were placed ad libitum on a control diet, fructose-free diet, or high-fructose diet (60% fructose) for two weeks prior to tumor implantation and for the duration of their lifespan.

### BONE MARROW CHIMERA GENERATION

CD45.1 mice were subjected to 1000 rads in once dose. After 4 hours, 5 million cells isolated from either WT or GLUT5-KO mice was injected retro-orbitally. Bone marrow cells were isolated, washed twice with 50mL of serum-free RPMI supplemented with 20mM HEPES and penicillin/streptomycin, counted, and resuspended in PBS at 50 x 106 cells/mL. 100uL cells (5×106 cells) were injected retro-orbitally. 6 weeks post-transplant, percent engraftment was measured via flow cytometry, and tumor implantation was performed 8 weeks post-implant.

One week prior to irradiation, recipient mice were given acidified water (pH 2.6) with 0.1mg/mL neomycin (Sigma; N112) and 0.01mg/mL polymyxin B sulphate (Sigma; P4932). Mice were kept on the acidified, antibiotic water for 2 weeks post-transplantation and then switched to acidic water for the duration of their lifespan.

### IMMUNOPHENOTYPING

Brains, spleens, and deep cervical lymphnodes were harvested from naïve or tumor-bearing mice into single-cell suspension. Cells were stained with BD Horizon™ Fixable Viability Stain 780 (BD Biosciences; 565388), blocked with anti-CD16/32, and stained with cell surface markers. Prior to intracellular staining, cells were resuspended in the eBioscience™ Foxp3 / Transcription Factor Staining Buffer Set (ThermoFisher; 00-5523-00) according to manufacturer’s instructions. For cytokine staining, cells were stimulated for 4h with eBioscience™ Cell Stimulation Cocktail (plus protein transport inhibitors) (500X) (ThermoFisher; 00-4975-93) prior to staining. Data were acquired using the BD FACSymphony™ A5 SE Cell Analyzer, the BD FACSymphony™ A5 Cell Analyzer, or the BD LSRFortessa™ Cell Analyzer. All colors were compensated with single-color controls from splenocytes.

### qRTPCR

RNA was extracted from cells using the Omega E.Z.N.A. RNA Isolation Kit (R6834-02, Omega Biol-tek, Norcross, GA) per manufacturer’s instructions. cDNA was generated using the iScript cDNA Synthesis Kit (BioRad; 1708891) and qPCR was performed using iTaq™ Universal SYBR® Green Supermix (Biorad; 172512) and rad on a CFX96 qPCR machine (Biorad). Primers were as follows: b-Actin (mouse) (forward 5’-TGGCAACTGTTCCTG-3’; reverse 5’-GGAAGCAGCCCTTCATCTTT-3’); Il1b (mouse) (forward 5’-CTAAGGCCAACCGTGAAAAG-3’; reverse 5’-ACCAGAGGCATACAGGGACA -3’); GLUT5 (mouse) (forward 5’-AAGCGACGACGTCCAGTATGT-3’; reverse 5’-GAATCGCCGTCCCCAAAG-3’)

### PRIMARY MICROGLIA ISOLATION AND CULTURE

Brains from P1-P4 neonatal mice were harvested and dissociated using the adult brain dissociation kit (Miltenyi Biotec 130-107-677) according to manufacturer’s instructions. Microglia were isolated using CD11b(Microglia) MicroBeads (Miltenyi Biotec; 130-093-634) according to manufacture’s instructions. Isolated microglia were resuspended in a previously established TIC media^68^ counted and spot plated in 50uL at 16 cells/mL. Cells were allowed to adhere for 30m on collagen-IV coated plates in standard tissue culture incubators before addition of 1mL TIC media containing either 5mM glucose or 5mM glucose and 5mM fructose. The next day, microglia were transferred to an incubator at 3.5% oxygen and cultured for 2-5 days before utilization for experimental assays.

### PHAGOCYTOSIS

CT-2A glioma cells were treated overnight with 50nM staurosporine (Caymen Chemical; 81590). Apoptotic cells were lifted with trypsin, spun down, and resuspended in warmed HBSS containing 1uM Cytiva CypHer5E NHS Esters (Fisher; 09-928-188). Staining was performed for 45m in a 37C water bath protected from light with gentle inversion every 10m. After 45m, cells were spun down, resuspended in complete DMEM, and incubated in water bath for 20m to remove and neutralize excess dye. Cells were then spun, resuspended in 1mL TIC media, and counted. Microglia were stained with 1uM CellTracker Green CMFDA (Invitrogen; C7025) for 20m in TIC media containing either 5mM glucose or 5mM glucose and 5mM fructose. Wells were washed 2x with complete DMEM before returning cells to indicated TIC media. Stained apoptotic CT-2A cells were added to wells at a ratio of 1:1 for and incubated at 3.5% O2 37C for 30 minutes. Cells were washed with cold FACS (PBS 2% FBS) buffer 1-2 times and gently scraped into 300uL FACS buffer before data was acquired via flow cytometry.

### RNASEQ

Microglia were isolated from adult mice using the adult brain dissociation kit (Miltenyi Biotec 130-107-677) and using CD11b(Microglia) MicroBeads (Miltenyi Biotec; 130-093-634) according to the manufacturer’s instructions. RNA was extracted using the RNeasy Plus Micro Kit (Qiagen; 74034). RNASequencing and analysis was performed by Novogene.

### scRNA-seq

CT-2A tumor cells were implanted intracranially, and the resulting tumor was removed after 14 days and digested into single-cell suspension using the adult brain dissociation kit (Miltenyi) according to the manufacturers instructions. CD45+ cells were isolated via magnetic bead isolation (Miltenyi) and submitted to the Northwestern University NUseq facility core for single cell library preparation and sequencing. AOPI fluorescent staining was used to determine cell number and viability Nexcelom Cellometer Auto2000 with. Sixteen thousand cells were loaded into the Chromium Controller (10X Genomics, PN-120223) on a Chromium Next GEM Chip K (10X Genomics, PN-1000127), and processed to generate single cell gel beads in the emulsion (GEM) according to the manufacturer’s protocol. The cDNA and library were generated using the Chromium Next GEM Single Cell 5’ Reagent Kits v2 (10X Genomics, PN-1000283) according to the manufacturer’s manual. In addition, mouse T-cell and B-cell V(D)J libraries were constructed using Chromium Single cell mouse TCR and BCR Amplification kit (10X Genomics, PN-1000252 and PN-1000255). The multiplexed libraries were pooled and sequenced on Illumina HiSeq 4000 sequencer with paired-end 50 kits using the following read length: 28 bp Read1 for cell barcode and UMI and 91 bp Read2 for transcript. The targeted sequencing depth for gene expression, mouse T-cell and B-cell V(D)J library is 20000, 5000, and 5000 reads per cell, respectively.

Raw sequencing files were aligned to mm10 reference using Cell Ranger (v3.1.0). The Seurat R Package using the scRNA-seq Seurat10x genomic workflow was used for all analyses unless noted otherwise [29608179]. Cells were filtered using a percent mitochondrial DNA threshold of 10% and a UMI range of 200 to 5000. Cells were then subject to Log Normalize, Scale Data, and PCA functions. The FindClusters and FindMarkers functions were utilized for clustering and marker identification and non-linear dimensional reduction techniques were applied to visual data in UMAP plot format. The Harmony algorithm was used to regress batch effects [31740819]. Immune cell clusters were annotated using three methods to produce robust cell assignments: 1) examination of DEGs against known lineage and functional markers^69–73^ singleR R package, an automated cell assignment algorithm^74^.

### TCR Analysis

TCR analysis was conducted using scRepertoire R package using standard workflow^75^. Clones combined between samples using combineTCR and then unique clones were found using cloneCall = “aa” and then merged into processed Seurat object for visualization and further analysis.

### GENE ONTOLOGY ENRICHMENT ANALYSIS

The DEGs were used for Gene Ontology enrichment analysis using the Bioconductor Package Cluster Profiler^76^. Significantly enriched GO-BP (Gene Ontology-Biological processes) terms were retrieved by setting the threshold of FDR=3; queried genes were manually selected using immunological keywords. Results were displayed using bubble plots, ridges and GSEA plots.

### INTERACTOME ANALYSIS

Potential ligand receptor (LR) interactions were analyzed using the Bioconductor package CellChat^77^. Only immune populations annotated as myeloid or lymphoid were included in the analysis. CellChat objects for BRAF-Fusion and NB were created, and a comparison analysis was used to infer differentially enriched ligand-receptor interactions between the tumor subtypes. This was performed using compareInteractions and RankNet functions. Results were displayed using heatmaps and circle diagrams to visualize significant interactions occurring within the TME.

### AUTOMATED MULTIPLEXED SEQUENTIAL IMMUNOFLUORESCENCE IMAGING

Automated hyperplex IF staining and imaging was performed on FFPE sections using the COMET™ system (Lunaphore Technologies). The multiplex panel included these antibodies: GLUT5 (Thermofisher, clone 14C8, cat#MA1-036X, dilution 1/100), P2RY12 (Atlas antibodies, cat#HPA014518, dilution 1/1000), CD31 (Abcam, clone EPR17259, cat#ab225883, dilution 1/1500), GFAP (Sigma, clone GA5, Cat#MAB360, dilution 1/3000), CD163 (Abcam, clone EPR19518, cat#ab182422, dilution 1/600), and CD68 (Dako Agilent, clone PG-M1, cat#GA613, dilution 1/1). The protocol was generated using the COMET™ Control Software, and reagents were loaded onto the COMET™ device to perform the seqIF™ protocol. All antibodies were validated using conventional IHC and/or IF staining in conjunction with corresponding fluorophores and 4’,6-diamidino-2-pheynlindole counterstain (DAPI, ThermoFisher Scientific). For optimal concentration and best signal-to-noise ratio, all antibodies were tested at 3 different dilutions, starting with the manufacturer-recommended dilution (MRD), MRD/2, and MRD/4. Secondary Alexa fluor 555 (ThermoFisher Scientific) and Alexa fluor 647 (ThermoFisher Scientific) were used at 1/200 and 1/400 dilutions, respectively. The optimizations and full runs of the multiplexed panel were executed using integrated protocols in the COMET™. All reagents were diluted in Multistaining Buffer (BU06, Lunaphore Technologies). Elution step lasted 2min for each cycle and was performed with Elution Buffer (BU07-L, Lunaphore Technologies) at 37°C. Quenching step lasted for 30sec and was performed with Quenching Buffer (BU08-L, Lunaphore Technologies). Imaging step was performed with Imaging Buffer (BU09, Lunaphore Technologies). Staining incubation times were set at 4min for all primary antibodies and at 2min secondary antibodies. Imaging step is performed with an integrated epifluorescent microscope at 20x magnification. Image registration was performed immediately after concluding the protocol by the control Software. Each seqIF™ protocol resulted in a multi-layer OME-TIFF file where the imaging outputs from each cycle are stitched and aligned. The OME-TIFF file contains DAPI image, intrinsic tissue autofluorescence in TRITC and Cy5 channels, and a single fluorescent layer per marker. Markers were subsequently pseudocolored for visualization of multiplexed antibodies using the HORIZON Viewer™ software. Analysis was performed using ImageJ.

### SEAHORSE XF96 EXTRACELLULAR FLUX ANALYSIS

An XFe86/XF Pro FluxPak (Agilent; 103022-100) cartridge was hydrated at 37C overnight in water in a non-CO2 incubator. Four hours before assay, the water was switched to XF Calibrant. BV2 microglia and RAW macrophages were lifted, washed with Seahorse XF Media (Agilent 103335-100) containing 2mM glutamine (Agilent 103579-100) and were seeded at 4×104 cells per well using CellTak (Corning; 354240) per manufacturer’s instructions. The seahorse assay was then run with a first injection of either 5mM glucose or 5mM fructose followed by the Seahorse XF Cell Mito Stress Test Kit (Agilent; 103015-100) per manufacturer’s instructions (Oligomycin 2uM; FCCP 1uM; Rotenone/Antimycin A 1uM). 25mM 2-DG (Sigma; D8375) was included in the final injection alongside Rotenone/Antimycin A.

### FRUCTOSE AND GLUCOSE DERIVATIZATION AND GAS CHROMATOGRAPHY-MASS SPECTROMETRY (GC-MS) ANALYSIS

5µL samples, glucose standards, and fructose standards were extracted with 30µL methanol containing 0.83mM 13C6-Glucose. Samples were vortexed, incubated on ice for 20 min, centrifuged at 21,000g at 4 °C for 20 min. 27.5uL of the supernatant was transferred to a glass GC–MS vial and dried down using nitrogen gas. Calibration standards were prepared from 7.81μM to 1mM for fructose and 31.25μM to 16mM for glucose and extracted along samples. We used a two-step derivatization procedure. First, methoxyamination was performed by adding 12.5μl of methoxyamine hydrochloride (20 mg/ml in pyrimidine) to the dried samples and incubating at 30 °C for 90 min. Then, silylation was carried out by adding 20μl of MSTFA (N-methyl-N-(trimethylsilyl)trifluoroacetamide) plus 1% TMCS (2,2,2-trifluoro-N-methyl-N-(trimethylsilyl)-acetamide, chlorotrimethylsilane) and incubating at 70°C for 60 min. Derivatized samples were cooled down to room temperature before injection.

Samples were analyzed by gas chromatography coupled with mass spectrometry using a 8890 gas chromatograph system (Agilent Technologies) with a HP-5ms Ultra Inert Column (Agilent Technologies, 19091S-433UI) coupled with an 5997B Mass Selective Detector (MSD). Helium was used as the carrier gas at a constant pressure of 14p.s.i. One microliter of sample was injected using a 1:3 split flow at 250°C. After injection, the GC oven was held at 60°C for 1min, increased to 320°C at 10°C per min, held at 320°C for 5min. The MS system operated under electron impact ionization at 70eV, and the MS source and quadrupole were held at 300°C. The detector was used in scanning mode with an ion range of 50–750m/z. Metabolites were analyzed using Masshunter software, with the following parameters. 12C6-Glucose: m/z = 319.2, RT = 16.9 and 17.1 min; 13C6-Glucose: m/z = 323.2 m/z, RT = 16.9 and 17.1 min; 12C6-Fructose: m/z = 217.2 m/z, RT = 16.6 and 16.75 min. 12C6-metabolite peak intensities were normalized to 13C6-Glucose peak intensity and the linear range of the calibration standards were used for absolute quantification of the metabolites.

### GSH/GSSG

Primary microglia were stimulated with 1ug/mL LPS for 4h in TIC media containing either 5mM glucose or 5mM glucose/5mM fructose. Assay was performed using the GSH/GSSG-Glo™ Assay (Promga; V6611) according to manufacturer’s instructions. GSH/GSSG ratios were calculated based on RLU and reported as the GSH/GSSG of glucose/fructose-cultured cells relative to the ratio in glucose alone cultured cells for that sample.

### QUANTIFICATION AND STATISTICAL ANALYSIS

Asside from RNAseq and scRNA-seq, all statistical analysis was performed using Prism software (Graphpad, Sand Diego, CA). Statistical significant between two groups was determined using two-tailed unpaired Student’s t test. Kaplan-Meier curves were generated, and log-rank test was performed to determine significance of in vivo survival rates. P values are indicated in the figures. Error bars are shown as ±SEMs for all figures.

---

**Extended Data 1.**
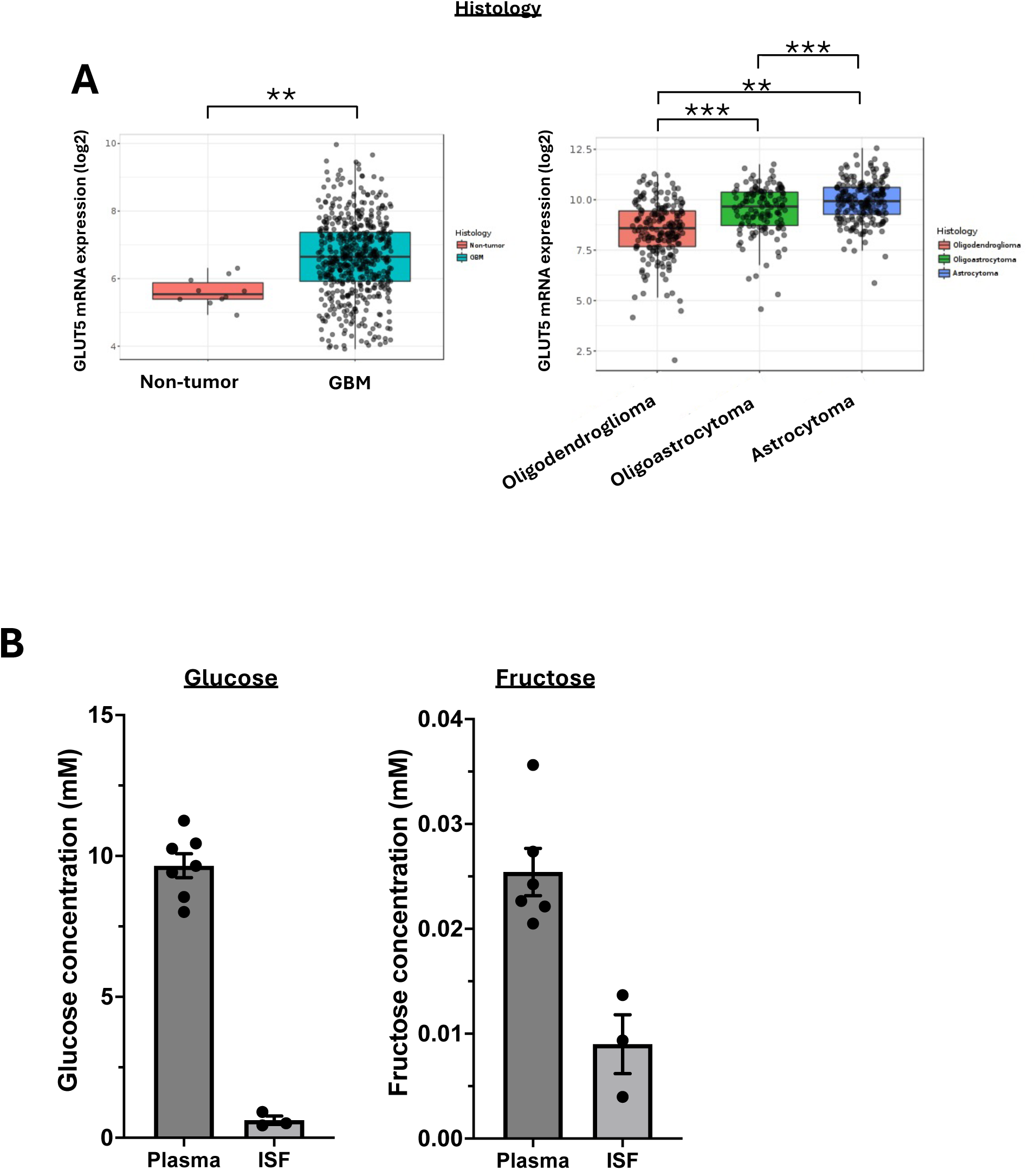
**A.** GLUT5 expression in non-tumor and GBM brain from human patients from GlioVis^78^. **B**. GLUT5 expression in oligodendroglioma, oligoastrocytoma, and astrocytoma from human patients from GlioVis^78^. **C**. Glucose concentration in mouse plasma and ISF (N=7 plasma; N=3 ISF). **D**. Fructose concentration in mouse plasma and ISF (N67 plasma; N=3 ISF)

**Extended Data 2.**
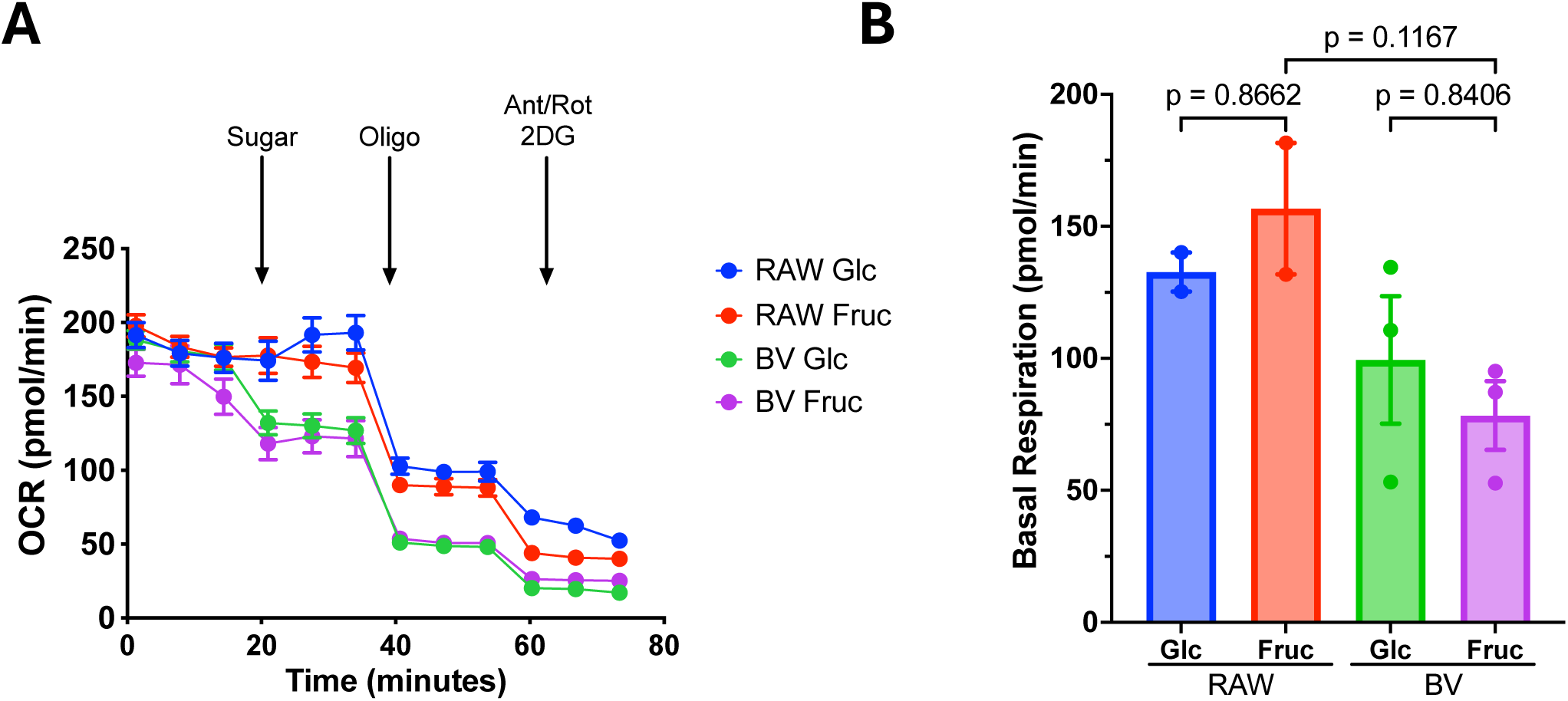
**A**. Basal respiration (oxygen consumption rate) of RAW or BV2 cells fed 5mM glucose or 5mM fructose (N=2 RAW, N=3 BV2; representative figure shown). **B**. Basal respiration in RAW or BV2 cells fed with 5mM glucose or 5mM fructose (N=2 RAW, N=3 BV2)

**Extended Data 3.**
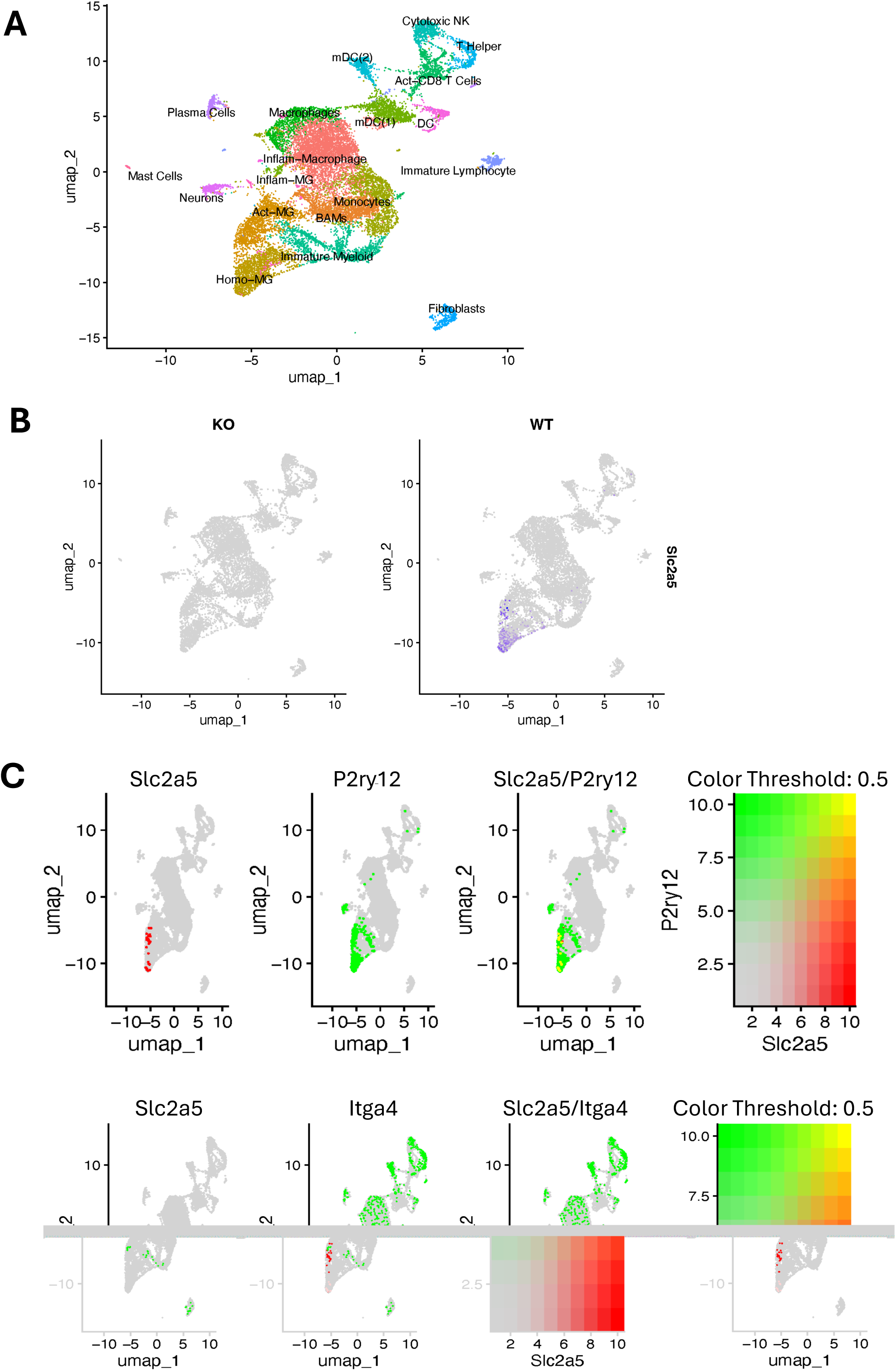
**A**. UMAP of immune cells from scRNA-sequencing of CD45+ cells isolated from tumors from C57Bl/6 mice implanted with 100K CT-2A glioma cells. **B**. UMAP of Slc2a5 expression from scRNA-sequencing of CD45+ cells isolated from tumors in WT or GLUT5 KO mice implanted with 100K CT-2A glioma cells. **C**. Normalized gene expression of Slc2a5 (left), P2ry12 (center) and Slc2a5/P2ry12 merged in immune cell UMAP in panel A. **D**. Normalized gene expression of Slc2a5 (left), Itga4 (center) and Slc2a5/P2ry12 merged in immune cell UMAP in panel A.

**Extended Figure 4.**
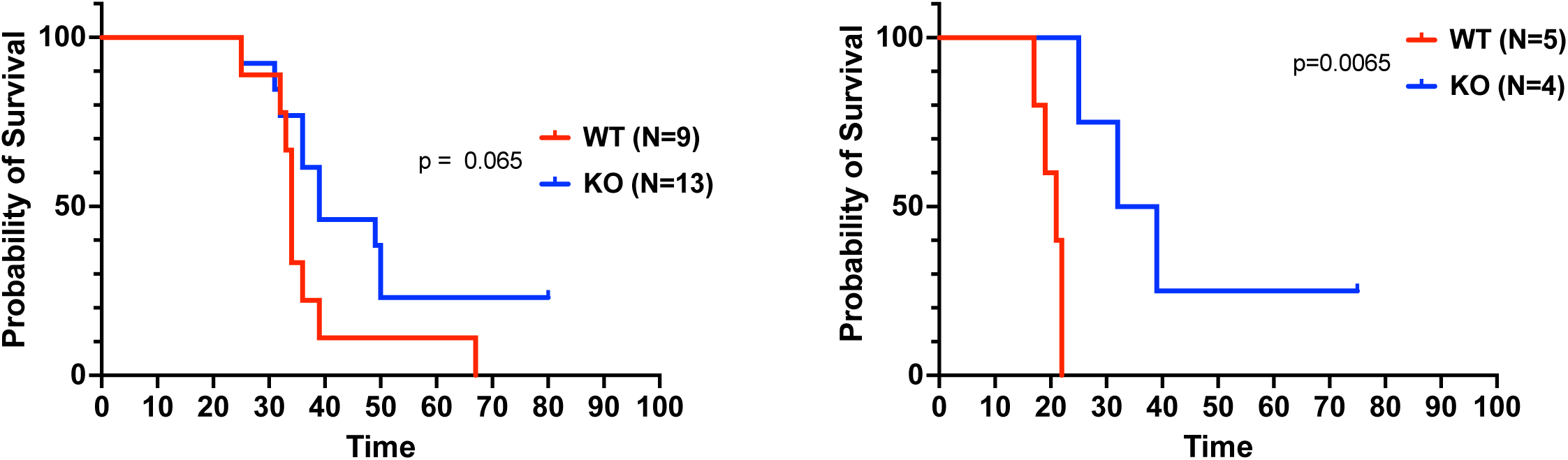
Kaplan-Meier curves of WT and GLUT5-KO mice implanted i.c. with 1×10^5 CT-2A glioma cells.

**Extended Figure 5.**
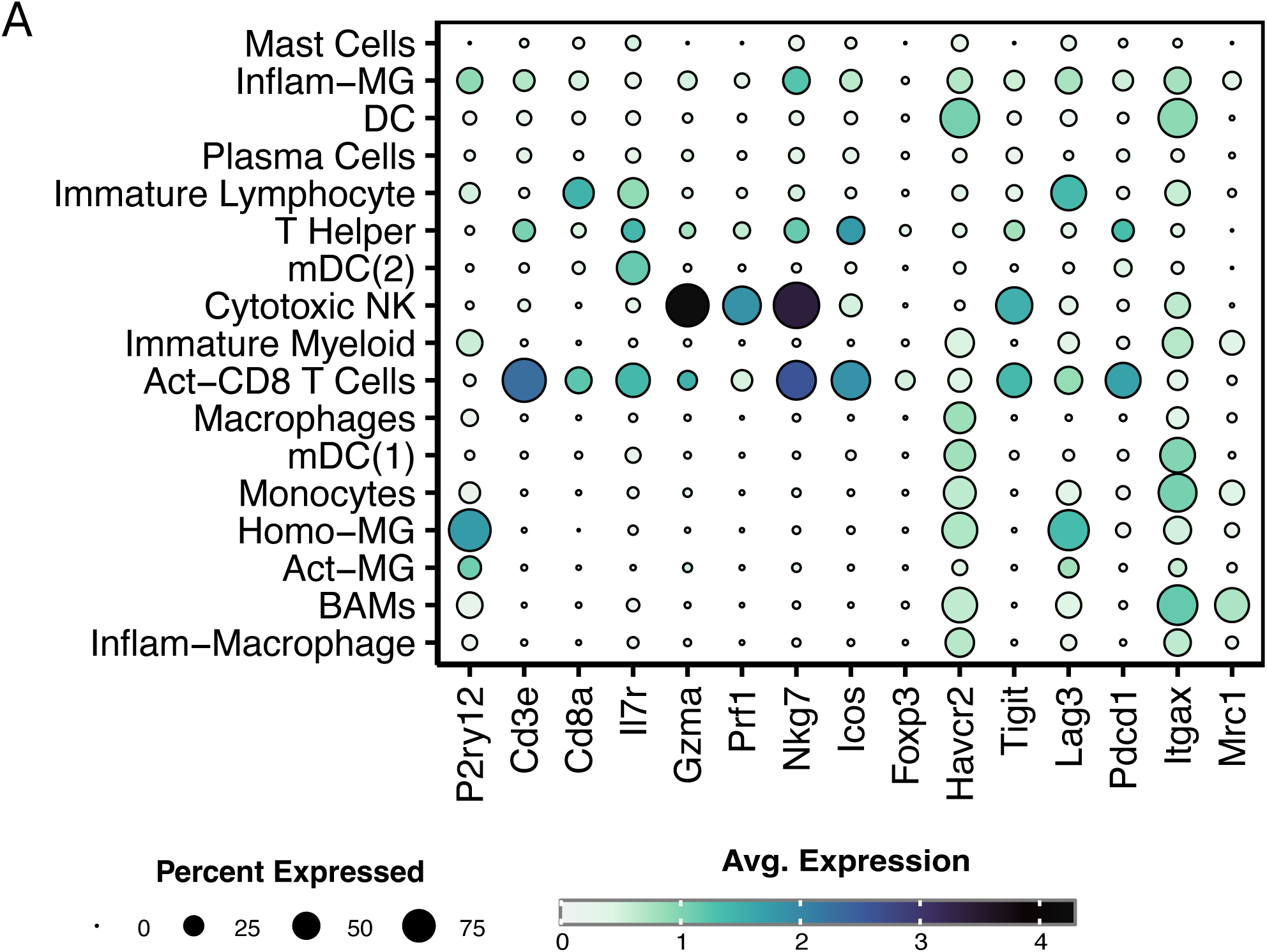
Identification and description of immune cell subsets in CD45+ cells in CT-2A glioma cell tumors

## REFERENCES

1. Stupp, R., Mason, W.P., van den Bent, M.J., Weller, M., Fisher, B., Taphoorn, M.J.B., Belanger, K., Brandes, A.A., Marosi, C., Bogdahn, U., et al. (2005). Radiotherapy plus concomitant and adjuvant temozolomide for glioblastoma. N Engl J Med 352, 987–996. 10.1056/NEJMoa043330.

2. Nduom, E.K., Weller, M., and Heimberger, A.B. (2015). Immunosuppressive mechanisms in glioblastoma. Neuro Oncol 17 *Suppl 7*, vii9–vii14. 10.1093/neuonc/nov151.

3. Thorsson, V., Gibbs, D.L., Brown, S.D., Wolf, D., Bortone, D.S., Ou Yang, T.-H., Porta-Pardo, E., Gao, G.F., Plaisier, C.L., Eddy, J.A., et al. (2018). The Immune Landscape of Cancer. Immunity 48, 812–830.e14. 10.1016/j.immuni.2018.03.023.

4. Zhang, P., Miska, J., Lee-Chang, C., Rashidi, A., Panek, W.K., An, S., Zannikou, M., Lopez-Rosas, A., Han, Y., Xiao, T., et al. (2019). Therapeutic targeting of tumor-associated myeloid cells synergizes with radiation therapy for glioblastoma. Proc Natl Acad Sci U S A 116, 23714–23723. 10.1073/pnas.1906346116.

5. Akkari, L., Bowman, R.L., Tessier, J., Klemm, F., Handgraaf, S.M., de Groot, M., Quail, D.F., Tillard, L., Gadiot, J., Huse, J.T., et al. (2020). Dynamic changes in glioma macrophage populations after radiotherapy reveal CSF-1R inhibition as a strategy to overcome resistance. Sci Transl Med 12, eaaw7843. 10.1126/scitranslmed.aaw7843.

6. Baghdadi, M., Wada, H., Nakanishi, S., Abe, H., Han, N., Putra, W.E., Endo, D., Watari, H., Sakuragi, N., Hida, Y., et al. (2016). Chemotherapy-Induced IL34 Enhances Immunosuppression by Tumor-Associated Macrophages and Mediates Survival of Chemoresistant Lung Cancer Cells. Cancer Res 76, 6030–6042. 10.1158/0008-5472.CAN-16-1170.

7. Liang, H., Deng, L., Hou, Y., Meng, X., Huang, X., Rao, E., Zheng, W., Mauceri, H., Mack, M., Xu, M., et al. (2017). Host STING-dependent MDSC mobilization drives extrinsic radiation resistance. Nat Commun 8, 1736. 10.1038/s41467-017-01566-5.

8. Kloosterman, D.J., Erbani, J., Boon, M., Farber, M., Handgraaf, S.M., Ando-Kuri, M., Sánchez-López, E., Fontein, B., Mertz, M., Nieuwland, M., et al. (2024). Macrophage-mediated myelin recycling fuels brain cancer malignancy. Cell 187, 5336–5356.e30. 10.1016/j.cell.2024.07.030.

9. Rashidi, A., Billingham, L.K., Zolp, A., Chia, T., Silvers, C., Katz, J.L., Park, C.H., Delay, S., Boland, L., Geng, Y., et al. (2024). Myeloid cell-derived creatine in the hypoxic niche promotes glioblastoma growth. Cell Metabolism 36, 62–77.e8. 10.1016/j.cmet.2023.11.013.

10. Miska, J., Rashidi, A., Lee-Chang, C., Gao, P., Lopez-Rosas, A., Zhang, P., Burga, R., Castro, B., Xiao, T., Han, Y., et al. (2021). Polyamines drive myeloid cell survival by buffering intracellular pH to promote immunosuppression in glioblastoma. Sci. Adv. 7, eabc8929. 10.1126/sciadv.abc8929.

11. Bernier, L.-P., York, E.M., Kamyabi, A., Choi, H.B., Weilinger, N.L., and MacVicar, B.A. (2020). Microglial metabolic flexibility supports immune surveillance of the brain parenchyma. Nat Commun 11, 1559. 10.1038/s41467-020-15267-z.

12. Payne, J., Maher, F., Simpson, I., Mattice, L., and Davies, P. (1997). Glucose transporter Glut 5 expression in microglial cells. Glia 21, 327–331. 10.1002/(sici)1098-1136(199711)21:3<327::aid-glia7>3.0.co;2-1.

13. Horikoshi, Y., Sasaki, A., Taguchi, N., Maeda, M., Tsukagoshi, H., Sato, K., and Yamaguchi, H. (2003). Human GLUT5 immunolabeling is useful for evaluating microglial status in neuropathological study using paraffin sections. Acta Neuropathol 105, 157–162. 10.1007/s00401-002-0627-4.

14. Altendorfer-Kroath, T., Hummer, J., and Birngruber, T. (2023). In vivo monitoring of brain pharmacokinetics and pharmacodynamics with cerebral open flow microperfusion. Biopharm Drug Dispos 44, 84–93. 10.1002/bdd.2343.

15. Hwang, J.J., Johnson, A., Cline, G., Belfort-DeAguiar, R., Snegovskikh, D., Khokhar, B., Han, C.S., and Sherwin, R.S. (2015). Fructose levels are markedly elevated in cerebrospinal fluid compared to plasma in pregnant women. PLoS One 10, e0128582. 10.1371/journal.pone.0128582.

16. Hannou, S.A., Haslam, D.E., McKeown, N.M., and Herman, M.A. (2018). Fructose metabolism and metabolic disease. J Clin Invest 128, 545–555. 10.1172/JCI96702.

17. Herman, M.A., and Birnbaum, M.J. (2021). Molecular aspects of fructose metabolism and metabolic disease. Cell Metab 33, 2329–2354. 10.1016/j.cmet.2021.09.010.

18. Wang, Z., Lipshutz, A., Liu, Z.-L., Trzeciak, A.J., Miranda, I.C., Martínez De La Torre, C., Schild, T., Lazarov, T., Rojas, W.S., Saavedra, P.H.V., et al. (2023). Early life high fructose exposure disrupts microglia function and impedes neurodevelopment. Preprint, 10.1101/2023.08.14.553242 https://doi.org/10.1101/2023.08.14.553242.

19. Altendorfer-Kroath, T., Asslaber, M., Hummer, J., Boulgaropoulos, B., Prietl, B., Pieber, T.R., Bernhart, E., and Birngruber, T. (2023). Atraumatic access to human glioblastoma in a xenograft animal model by cerebral open flow microperfusion. J Neurosci Methods 393, 109893. 10.1016/j.jneumeth.2023.109893.

20. Sullivan, M.R., Lewis, C.A., and Muir, A. (2019). Isolation and Quantification of Metabolite Levels in Murine Tumor Interstitial Fluid by LC/MS. Bio Protoc 9, e3427. 10.21769/BioProtoc.3427.

21. Sullivan, M.R., Danai, L.V., Lewis, C.A., Chan, S.H., Gui, D.Y., Kunchok, T., Dennstedt, E.A., Vander Heiden, M.G., and Muir, A. (2019). Quantification of microenvironmental metabolites in murine cancers reveals determinants of tumor nutrient availability. Elife 8, e44235. 10.7554/eLife.44235.

22. Wishart, D.S., Guo, A., Oler, E., Wang, F., Anjum, A., Peters, H., Dizon, R., Sayeeda, Z., Tian, S., Lee, B.L., et al. (2022). HMDB 5.0: the Human Metabolome Database for 2022. Nucleic Acids Res 50, D622–D631. 10.1093/nar/gkab1062.

23. Buchs, A.E., Sasson, S., Joost, H.G., and Cerasi, E. (1998). Characterization of GLUT5 domains responsible for fructose transport. Endocrinology 139, 827–831. 10.1210/endo.139.3.5780.

24. Hundal, H.S., Darakhshan, F., Kristiansen, S., Blakemore, S.J., and Richter, E.A. (1998). GLUT5 Expression and Fructose Transport in Human Skeletal Muscle. In Skeletal Muscle Metabolism in Exercise and Diabetes Advances in Experimental Medicine and Biology., E. A. Richter, B. Kiens, H. Galbo, and B. Saltin, eds. (Springer US), pp. 35–45. 10.1007/978-1-4899-1928-1_4.

25. Concha, I.I., Velásquez, F.V., Martínez, J.M., Angulo, C., Droppelmann, A., Reyes, A.M., Slebe, J.C., Vera, J.C., and Golde, D.W. (1997). Human erythrocytes express GLUT5 and transport fructose. Blood 89, 4190–4195.

26. Mueckler, M., and Thorens, B. (2013). The SLC2 (GLUT) family of membrane transporters. Mol Aspects Med 34, 121–138. 10.1016/j.mam.2012.07.001.

27. Karlsson, M., Zhang, C., Méar, L., Zhong, W., Digre, A., Katona, B., Sjöstedt, E., Butler, L., Odeberg, J., Dusart, P., et al. (2021). A single-cell type transcriptomics map of human tissues. Sci Adv 7, eabh2169. 10.1126/sciadv.abh2169.

28. Single cell type - SLC2A5 - The Human Protein Atlas https://www.proteinatlas.org/ENSG00000142583-SLC2A5/single+cell.

29. SLC2A5 protein expression summary - The Human Protein Atlas https://www.proteinatlas.org/ENSG00000142583-SLC2A5.

30. The Human Protein Atlas https://www.proteinatlas.org/.

31. Single cell type - SLC2A5 - The Human Protein Atlas.

32. SLC2A5 protein expression summary - The Human Protein Atlas.

33. The Human Protein Atlas.

34. Douard, V., and Ferraris, R.P. (2008). Regulation of the fructose transporter GLUT5 in health and disease. Am J Physiol Endocrinol Metab 295, E227–237. 10.1152/ajpendo.90245.2008.

35. Shay, T., and Kang, J. (2013). Immunological Genome Project and systems immunology. Trends Immunol 34, 602–609. 10.1016/j.it.2013.03.004.

36. Ruiz-Moreno, C., Salas, S.M., Samuelsson, E., Brandner, S., Kranendonk, M.E.G., Nilsson, M., and Stunnenberg, H.G. (2022). Harmonized single-cell landscape, intercellular crosstalk and tumor architecture of glioblastoma. Preprint, 10.1101/2022.08.27.505439 https://doi.org/10.1101/2022.08.27.505439.

37. Ravi, V.M., Neidert, N., Will, P., Joseph, K., Maier, J.P., Kückelhaus, J., Vollmer, L., Goeldner, J.M., Behringer, S.P., Scherer, F., et al. (2022). T-cell dysfunction in the glioblastoma microenvironment is mediated by myeloid cells releasing interleukin-10. Nat Commun 13, 925. 10.1038/s41467-022-28523-1.

38. Friebel, E., Kapolou, K., Unger, S., Núñez, N.G., Utz, S., Rushing, E.J., Regli, L., Weller, M., Greter, M., Tugues, S., et al. (2020). Single-Cell Mapping of Human Brain Cancer Reveals Tumor-Specific Instruction of Tissue-Invading Leukocytes. Cell 181, 1626–1642.e20. 10.1016/j.cell.2020.04.055.

39. Fixsen, B.R., Han, C.Z., Zhou, Y., Spann, N.J., Saisan, P., Shen, Z., Balak, C., Sakai, M., Cobo, I., Holtman, I.R., et al. (2023). SALL1 enforces microglia-specific DNA binding and function of SMADs to establish microglia identity. Nat Immunol 24, 1188–1199. 10.1038/s41590-023-01528-8.

40. Lund, H., Pieber, M., Parsa, R., Han, J., Grommisch, D., Ewing, E., Kular, L., Needhamsen, M., Espinosa, A., Nilsson, E., et al. (2018). Competitive repopulation of an empty microglial niche yields functionally distinct subsets of microglia-like cells. Nat Commun 9, 4845. 10.1038/s41467-018-07295-7.

41. Iorgulescu, J.B., Ruthen, N., Ahn, R., Panagioti, E., Gokhale, P.C., Neagu, M., Speranza, M.C., Eschle, B.K., Soroko, K.M., Piranlioglu, R., et al. (2023). Antigen presentation deficiency, mesenchymal differentiation, and resistance to immunotherapy in the murine syngeneic CT2A tumor model. Front Immunol 14, 1297932. 10.3389/fimmu.2023.1297932.

42. Chen, C.-H., Chin, R.L., Hartley, G.P., Lea, S.T., Engel, B.J., Hsieh, C.-E., Prasad, R., Roszik, J., Shingu, T., Lizee, G.A., et al. (2023). Novel murine glioblastoma models that reflect the immunotherapy resistance profile of a human disease. Neuro Oncol 25, 1415–1427. 10.1093/neuonc/noad025.

43. Shingu, T., Ho, A.L., Yuan, L., Zhou, X., Dai, C., Zheng, S., Wang, Q., Zhong, Y., Chang, Q., Horner, J.W., et al. (2017). Qki deficiency maintains stemness of glioma stem cells in suboptimal environment by downregulating endolysosomal degradation. Nat Genet 49, 75–86. 10.1038/ng.3711.

44. Gerrits, E., Heng, Y., Boddeke, E.W.G.M., and Eggen, B.J.L. (2020). Transcriptional profiling of microglia; current state of the art and future perspectives. Glia 68, 740–755. 10.1002/glia.23767.

45. DePaula-Silva, A.B., Gorbea, C., Doty, D.J., Libbey, J.E., Sanchez, J.M.S., Hanak, T.J., Cazalla, D., and Fujinami, R.S. (2019). Differential transcriptional profiles identify microglial- and macrophage-specific gene markers expressed during virus-induced neuroinflammation. J Neuroinflammation 16, 152. 10.1186/s12974-019-1545-x.

46. Barone, S., Fussell, S.L., Singh, A.K., Lucas, F., Xu, J., Kim, C., Wu, X., Yu, Y., Amlal, H., Seidler, U., et al. (2009). Slc2a5 (Glut5) is essential for the absorption of fructose in the intestine and generation of fructose-induced hypertension. J Biol Chem 284, 5056– 5066. 10.1074/jbc.M808128200.

47. Dimitrov, D., Türei, D., Garrido-Rodriguez, M., Burmedi, P.L., Nagai, J.S., Boys, C., Ramirez Flores, R.O., Kim, H., Szalai, B., Costa, I.G., et al. (2022). Comparison of methods and resources for cell-cell communication inference from single-cell RNA-Seq data. Nat Commun 13, 3224. 10.1038/s41467-022-30755-0.

48. Liu, H., Yan, Z., Zhu, D., Xu, H., Liu, F., Chen, T., Zhang, H., Zheng, Y., Liu, B., Zhang, L., et al. (2023). CD-NTase family member MB21D2 promotes cGAS-mediated antiviral and antitumor immunity. Cell Death Differ 30, 992–1004. 10.1038/s41418-023-01116-1.

49. Zhang, P., Rashidi, A., Zhao, J., Silvers, C., Wang, H., Castro, B., Ellingwood, A., Han, Y., Lopez-Rosas, A., Zannikou, M., et al. (2023). STING agonist-loaded, CD47/PD-L1-targeting nanoparticles potentiate antitumor immunity and radiotherapy for glioblastoma. Nat Commun 14, 1610. 10.1038/s41467-023-37328-9.

50. Yan, H., Wang, Z., Teng, D., Chen, X., Zhu, Z., Chen, H., Wang, W., Wei, Z., Wu, Z., Chai, Q., et al. (2024). Hexokinase 2 senses fructose in tumor-associated macrophages to promote colorectal cancer growth. Cell Metab 36, 2449–2467.e6. 10.1016/j.cmet.2024.10.002.

51. Park, J., Min, J.-S., Kim, B., Chae, U.-B., Yun, J.W., Choi, M.-S., Kong, I.-K., Chang, K.-T., and Lee, D.-S. (2015). Mitochondrial ROS govern the LPS-induced pro-inflammatory response in microglia cells by regulating MAPK and NF-κB pathways. Neurosci Lett 584, 191–196. 10.1016/j.neulet.2014.10.016.

52. Morris, G., Gevezova, M., Sarafian, V., and Maes, M. (2022). Redox regulation of the immune response. Cell Mol Immunol 19, 1079–1101. 10.1038/s41423-022-00902-0.

53. Canton, M., Sánchez-Rodríguez, R., Spera, I., Venegas, F.C., Favia, M., Viola, A., and Castegna, A. (2021). Reactive Oxygen Species in Macrophages: Sources and Targets. Front Immunol 12, 734229. 10.3389/fimmu.2021.734229.

54. Russell, D.G., Huang, L., and VanderVen, B.C. (2019). Immunometabolism at the interface between macrophages and pathogens. Nat Rev Immunol 19, 291–304. 10.1038/s41577-019-0124-9.

55. Simpson, D.S.A., and Oliver, P.L. (2020). ROS Generation in Microglia: Understanding Oxidative Stress and Inflammation in Neurodegenerative Disease. Antioxidants (Basel) 9, 743. 10.3390/antiox9080743.

56. Yan, L.-J. (2018). Redox imbalance stress in diabetes mellitus: Role of the polyol pathway. Animal Model Exp Med 1, 7–13. 10.1002/ame2.12001.

57. Schwaller, M., Wilkinson, B., and Gilbert, H.F. (2003). Reduction-reoxidation cycles contribute to catalysis of disulfide isomerization by protein-disulfide isomerase. J Biol Chem 278, 7154–7159. 10.1074/jbc.M211036200.

58. Todoric, J., Di Caro, G., Reibe, S., Henstridge, D.C., Green, C.R., Vrbanac, A., Ceteci, F., Conche, C., McNulty, R., Shalapour, S., et al. (2020). Fructose stimulated de novo lipogenesis is promoted by inflammation. Nat Metab 2, 1034–1045. 10.1038/s42255-020-0261-2.

59. Jaiswal, N., Agrawal, S., and Agrawal, A. (2019). High fructose-induced metabolic changes enhance inflammation in human dendritic cells. Clinical and Experimental Immunology 197, 237–249. 10.1111/cei.13299.

60. Ting, K.K.Y. (2024). Fructose-induced metabolic reprogramming of cancer cells. Front Immunol 15, 1375461. 10.3389/fimmu.2024.1375461.

61. Hwang, J.J., Jiang, L., Hamza, M., Dai, F., Belfort-DeAguiar, R., Cline, G., Rothman, D.L., Mason, G., and Sherwin, R.S. (2017). The human brain produces fructose from glucose. JCI Insight 2, e90508. 10.1172/jci.insight.90508.

62. Xu, J., Begley, P., Church, S.J., Patassini, S., McHarg, S., Kureishy, N., Hollywood, K.A., Waldvogel, H.J., Liu, H., Zhang, S., et al. (2016). Elevation of brain glucose and polyol-pathway intermediates with accompanying brain-copper deficiency in patients with Alzheimer’s disease: metabolic basis for dementia. Sci Rep 6, 27524. 10.1038/srep27524.

63. Kang, Y.-L., Kim, J., Kwak, S.-B., Kim, Y.-S., Huh, J., and Park, J.-W. (2024). The polyol pathway and nuclear ketohexokinase A signaling drive hyperglycemia-induced metastasis of gastric cancer. Exp Mol Med 56, 220–234. 10.1038/s12276-023-01153-3.

64. Schwab, A., Siddiqui, M.A., Ramesh, V., Gollavilli, P.N., Turtos, A.M., Møller, S.S., Pinna, L., Havelund, J.F., Rømer, A.M.A., Ersan, P.G., et al. (2024). Polyol pathway-generated fructose is indispensable for growth and survival of non-small cell lung cancer. Cell Death Differ. 10.1038/s41418-024-01415-1.

65. Sasaki, A., Horikoshi, Y., Yokoo, H., Nakazato, Y., and Yamaguchi, H. (2003). Antiserum against human glucose transporter 5 is highly specific for microglia among cells of the mononuclear phagocyte system. Neurosci Lett 338, 17–20. 10.1016/s0304-3940(02)01332-0.

66. Schild, T., Wallisch, P., Zhao, Y., Wang, Y.-T., Haughton, L., Chirayil, R., Pierpont, K., Chen, K., Nunes-Violante, S., Cross, J., et al. (2025). Metabolic engineering to facilitate anti-tumor immunity. Cancer Cell 43, 552–562.e9. 10.1016/j.ccell.2025.02.004.

67. Hamed, A.A., Hua, K., Trinh, Q.M., Simons, B.D., Marioni, J.C., Stein, L.D., and Dirks, P.B. (2025). Gliomagenesis mimics an injury response orchestrated by neural crest-like cells. Nature 638, 499–509. 10.1038/s41586-024-08356-2.

68. Bohlen, C.J., Bennett, F.C., Tucker, A.F., Collins, H.Y., Mulinyawe, S.B., and Barres, B.A. (2017). Diverse Requirements for Microglial Survival, Specification, and Function Revealed by Defined-Medium Cultures. Neuron 94, 759–773.e8. 10.1016/j.neuron.2017.04.043.

69. Cheng, D., Qiu, K., Rao, Y., Mao, M., Li, L., Wang, Y., Song, Y., Chen, J., Yi, X., Shao, X., et al. (2023). Proliferative exhausted CD8+ T cells exacerbate long-lasting anti-tumor effects in human papillomavirus-positive head and neck squamous cell carcinoma. Elife 12, e82705. 10.7554/eLife.82705.

70. Ma, R.-Y., Black, A., and Qian, B.-Z. (2022). Macrophage diversity in cancer revisited in the era of single-cell omics. Trends Immunol 43, 546–563. 10.1016/j.it.2022.04.008.

71. Paolicelli, R.C., Sierra, A., Stevens, B., Tremblay, M.-E., Aguzzi, A., Ajami, B., Amit, I., Audinat, E., Bechmann, I., Bennett, M., et al. (2022). Microglia states and nomenclature: A field at its crossroads. Neuron 110, 3458–3483. 10.1016/j.neuron.2022.10.020.

72. Wang, R., Peng, L., Xiao, Y., Zhou, Q., Wang, Z., Tang, L., Xiao, H., Yang, K., Liu, H., and Li, L. (2023). Single-cell RNA sequencing reveals changes in glioma-associated macrophage polarization and cellular states of malignant gliomas with high AQP4 expression. Cancer Gene Ther 30, 716–726. 10.1038/s41417-022-00582-y.

73. Tripathi, S., Najem, H., Dussold, C., Pacheco, S., Du, R., Sooreshjani, M., Hurley, L., Chandler, J.P., Stupp, R., Sonabend, A.M., et al. (2024). Pediatric glioma immune profiling identifies TIM3 as a therapeutic target in BRAF fusion pilocytic astrocytoma. J Clin Invest 134, e177413. 10.1172/JCI177413.

74. Aran, D., Looney, A.P., Liu, L., Wu, E., Fong, V., Hsu, A., Chak, S., Naikawadi, R.P., Wolters, P.J., Abate, A.R., et al. (2019). Reference-based analysis of lung single-cell sequencing reveals a transitional profibrotic macrophage. Nat Immunol 20, 163–172. 10.1038/s41590-018-0276-y.

75. Borcherding, N., Bormann, N.L., and Kraus, G. (2020). scRepertoire: An R-based toolkit for single-cell immune receptor analysis. F1000Res 9, 47. 10.12688/f1000research.22139.2.

76. Yu, G., Wang, L.-G., Han, Y., and He, Q.-Y. (2012). clusterProfiler: an R package for comparing biological themes among gene clusters. OMICS 16, 284–287. 10.1089/omi.2011.0118.

77. Jin, S., Guerrero-Juarez, C.F., Zhang, L., Chang, I., Ramos, R., Kuan, C.-H., Myung, P., Plikus, M.V., and Nie, Q. (2021). Inference and analysis of cell-cell communication using CellChat. Nat Commun 12, 1088. 10.1038/s41467-021-21246-9.

78. Bowman, R.L., Wang, Q., Carro, A., Verhaak, R.G.W., and Squatrito, M. (2017). GlioVis data portal for visualization and analysis of brain tumor expression datasets. Neuro Oncol 19, 139–141. 10.1093/neuonc/now247.

